# Dynamic Antigen Expression and Intrinsic CTL Resistance in HIV Reservoir Clones

**DOI:** 10.64898/2026.02.22.707300

**Authors:** Isabella A. T. M. Ferreira, Alberto Herrera, Tan Thinh Huynh, Nathan L. Board, Emily Stone, Noemi L. Linden, Yanqin Ren, Cintia Bittar, Virender K. Pal, Shane Vedova, Ethan Naing, Parul Sinha, Ali Danesh, Cristian Ovies, Fitty Liu, Louise Leyre, Marie Canis, Ana Rafaela Teixeira, Susan Moir, Tae-Wook Chun, Colin Kovacs, Paul Zumbo, Doron Betel, Elias K. Halvas, Guinevere Q. Lee, Marina Caskey, Paul D. Bieniasz, Michel C. Nussenzweig, R. Brad Jones

**Author notes:** These authors contributed equally (Isabella A. T. M. Ferreira, Alberto Herrera).

## Abstract

Reservoirs of clonally expanded CD4^+^ T-cells harboring rebound-competent HIV proviruses persist lifelong during ART. Latency is considered the principal barrier to viral eradication and has resisted pharmacological reversal, yet it appears that sustained immune pressure may still erode reservoirs. Recent advances have yielded glimpses into these exceptionally rare reservoir-harboring cells, implicating intrinsic pro-survival properties in their persistence. Here, we isolate and characterize populations of authentic reservoir clones (ARCs) that robustly proliferate and accumulate while producing infectious virus, without overtly succumbing to viral cytopathic effects. At any given moment, only small fractions of ARCs expressed HIV proteins, a state remarkably unperturbed by potent TCR or mitogenic stimulation. Nevertheless, sustained co-culture with cytotoxic T-lymphocytes (CTL) revealed extensive time-integrated antigenic vulnerability, culling clonal expansion of some ARCs by >90%. Notably, a regulatory T-cell ARC displayed pronounced cell-intrinsic resistance to CTL – a longstanding hypothesis we now directly demonstrate – linked to low oxidative stress and reversed with desferoxamine, a hypoxic stress inducer and FDA-approved therapeutic. Overall, we provide novel insights into the vulnerabilities of reservoir clones to potent, sustained CTL pressure and highlight intrinsic resistance pathways as actionable therapeutic targets, opening opportunities for advancing immune-based HIV cure strategies.

## Introduction

HIV persists for life in individuals treated with antiretroviral therapy (ART), in a reservoir comprised primarily of clonally expanded CD4^+^ T-cells harboring integrated proviruses, which rapidly fuel viral rebound if ART is interrupted^1–5^. Efforts to engage the immune system to eliminate these reservoirs have focused on reversing HIV latency, but have shown little success – thought to be due to the insensitivity of bona fide reservoirs to reactivation, in contrast to models of latency^6–10^. Somewhat paradoxically, however, for a model of entrenched latency, virologic and immunologic measures speak to ongoing immunosurveillance during ART: HIV-specific T-cells persist and show signs of recent antigen encounter^11–13^, and reservoirs exhibit footprints of immune selection – elimination of intact versus defective proviruses^14,15^. Residual reservoir cells also become increasingly clonal and enriched for HIV integrations in transcriptionally unfavorable genomic regions, such as heterochromatin and zinc-finger (ZNF) genes^16–19^. Some clones, however, continue to transcribe HIV - in some cases contributing to detectable viremia^20,21^ - and can present features that raise the possibility that they are intrinsically resistant to elimination by cytotoxic T-cells (CTL) or natural killer (NK) cells^22–24^. However, the extreme scarcity of reservoir-harboring cells (1-10 per million CD4^+^ T-cells) has precluded direct testing of mechanisms that might explain their persistence as well as limited characterization of the dynamic interplay between latency, proliferation, and immune surveillance.

Here, we describe a platform for the isolation, expansion, and characterization of authentic reservoir clones (ARCs) from ART-suppressed individuals, enabling their investigation – validated by stable multiomic characteristics preserved over months in cell culture. Our studies of ARCs reveal the limited efficacy of ‘shocking’ provirus out of latency, yet show that ARCs display antigenic vulnerability to sustained CTL pressure. Further, we directly observe cell-intrinsic resistance to CTL in a highly proliferative ARC revealing a therapeutically reversible barrier to elimination.

### Authentic reservoir clones can be isolated and expanded

HIV is generally cytopathic to CD4^+^ T-cells, resulting in a very short half-life (∼1 day *in vivo*) for productively infected cells^25,26^. Similarly, *in vitro* infection of activated CD4^+^ T-cells results in extensive cell death. However, recent studies indicate that certain CD4^+^ T-cell clones harboring intact HIV proviruses, which have expanded *in vivo*, can survive proliferation *in vitro* following antigen stimulation^4,27–35^. By sorting for TCR-Vβ segments known to contain a prominent reservoir clone in a given donor, we previously generated T-cell lines enriched for CD4^+^ T-cells carrying replication-competent proviruses^35,36^. Leveraging these insights, we present approaches to isolate and expand pure populations of ‘authentic reservoir clones’ (ARCs) with or without pre-enrichment.

We employed two complementary approaches to isolate ARCs: following enrichment based on TCR-Vβ segments^35^, and unbiased limiting-dilution without pre-enrichment (**Fig. 1a**). Cultures were screened by ELISA for HIV-Gag production under antiretroviral suppression, ensuring detected virus resulted solely from proliferation of pre-existing infected cells, rather than from rounds of viral replication. Purity was initially assessed by droplet digital PCR (ddPCR), where a provirus-to-host gene (RPP30) ratio of ∼1:2 suggested a pure clone (**Fig. 1b**). This was subsequently confirmed through the detection of a single, unique proviral integration site and TCR RNA sequence. We isolated 7 ARCs, each expanding into populations ranging from millions to hundreds of millions of cells (**Fig. 1c**). Near full-length proviral sequencing revealed that 6 of these ARCs contained intact HIV proviruses, while one contained a deletion in the 5’-leader sequence (**Extended data Fig. 1**). The 2 ARCs isolated by TCR-Vβ segments correspond to the provirus-enriched T-cell lines that were studied in our preceding work^35,36^.

**Figure 1.**
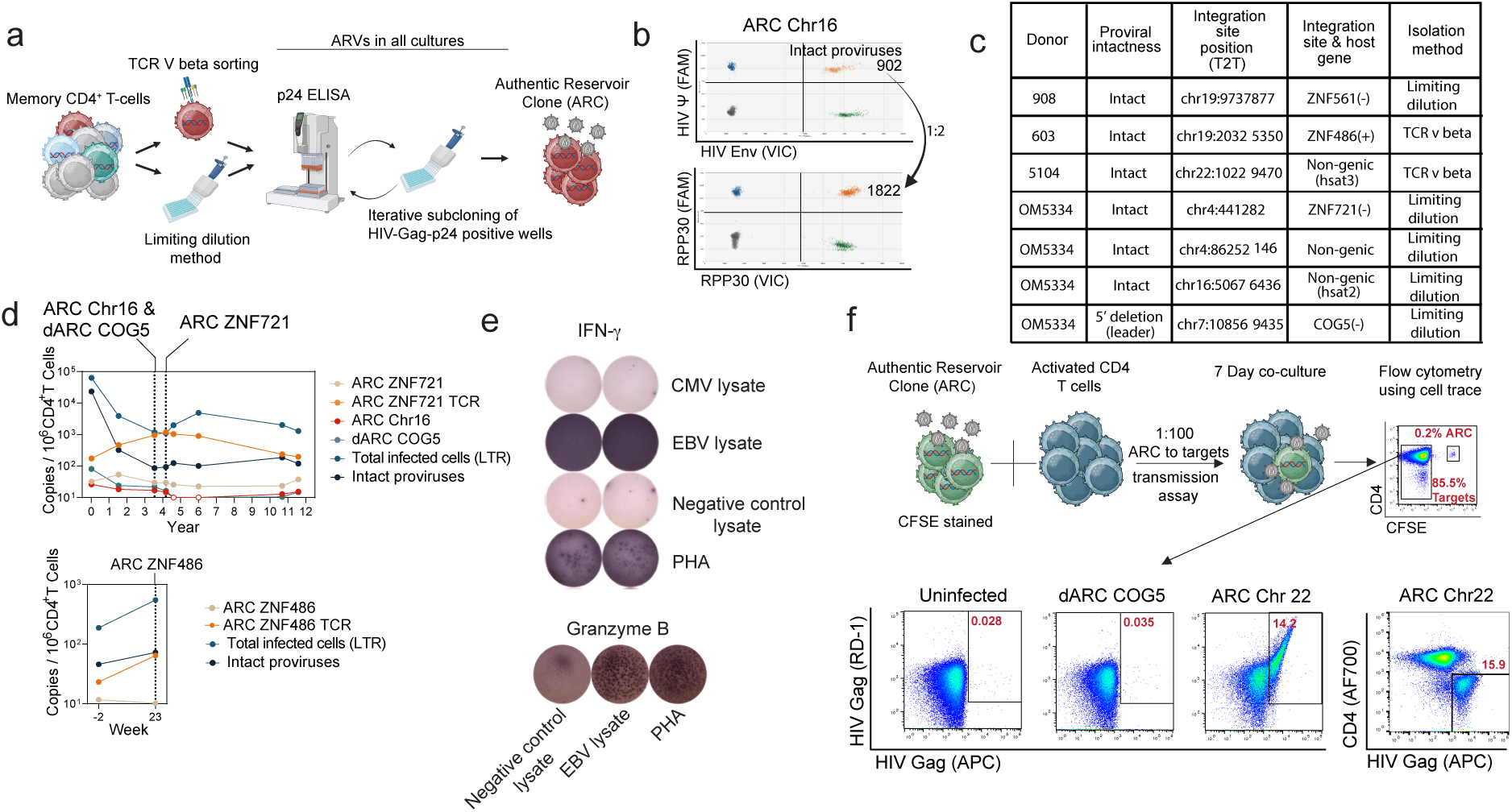
Isolation and characterization of authentic reservoir clones (ARC) **(a**) Schematic depicting isolation of ARCs via sorting for TCR Vβ associated with prominent reservoir clones, or by limiting dilution. Pure clones were derived by iterative subcloning of HIV-Gag-p24^+^ wells. **(b)** Intact Proviral DNA Assay (IPDA) confirming purity of isolated ARCs, indicated by a 1:2 provirus-to-RPP30 ratio. **(c)** Table summarizing six intact ARCs and one defective ARC (dARC) isolated from participants. (+) or (-) indicate sense or antisense orientation of integrated provirus. **(d)** Longitudinal analysis showing persistent detection of ARC ZNF721 over years on ART, quantified by TCR Vβ- and integration site-specific ddPCR. ARC ZNF486 was detected at the two timepoints available (labeled in relation to the timing of a bNAb intervention). ddPCR quantitation of total HIV LTRs and intact HIV provirus (IPDA) is provided for comparison. Dashed lines indicate timepoints from which the ARCs were isolated. **(e)** ELISPOT demonstrating EBV-specific IFN-γ and Granzyme B responses by ARC ZNF721. **(f)** Flow cytometry analysis showing transmission of replication-competent HIV from CFSE-labeled ARC Chr 22 to activated, CFSE-negative bulk CD4^+^ T-cells after 7-day co-culture without ARVs. For ARC Chr22, the infection is shown in two ways: with a dual Gag stain and depicting CD4 downregulation in HIV-expressing cells. As expected, no transmission of infectious virus occurs from the dARC.

### ARCs exhibit integration-site selection, stable persistence, and HIV rebound potential

HIV integrates quasi-randomly into the genome, with a preference for actively transcribed genes within accessible chromatin, but with proviruses broadly distributed across the genome^37,38^. However, following sustained immunologic pressure, clones with intact proviruses preferentially harbor integrations within specific genomic regions, notably zinc-finger (ZNF) genes or heterochromatic regions, while defectives including 5’-L defective proviruses, persist in gene bodies^16–19,39,40^. Strikingly, all 6 of the intact-provirus ARCs we isolated have integration sites in one of these regions – including the 4 isolated with the unbiased limiting dilution method (**Fig. 1c**). In contrast, the 5’-L defective provirus was integrated into the gene *COG5*. These findings suggest a concordance between the clones that are isolated and expanded in our system and those that survive sustained immune pressure *in vivo,* providing an opportunity to study their mechanisms of persistence (**Fig. 1c**).

Three of the ARCs with intact proviruses as well as the ARC with the defective provirus (termed a dARC) were isolated from a donor (OM5334, **Supplementary Table 1**) with samples spanning nearly 12 years, beginning approximately 4 months after HIV infection and within a week of ART initiation. Using ARC-specific sequencing data, we designed custom ddPCR assays to track total clonal populations (infected and uninfected) via rearranged TCR sequences and specifically infected subsets via proviral integration sites in each donors’ *ex vivo* samples to quantify the abundance of the parent cell (both infected and uninfected). These were complemented by standard assays for quantifying total HIV-infected cells (LTR) and intact HIV proviruses (IPDA). Each ARC comprised a small fraction of infected cells at ART initiation but showed remarkable stability over the subsequent 12 years (**Fig. 1d**), contrasting sharply with substantial declines in overall intact proviruses over the first 3.5 years. This long-term persistence demonstrates that these ARCs are substantially more resilient to the selective pressures present in this donor than most HIV-infected cells.

‘ARC ZNF721’ named for its proviral integration site in the *ZNF721* gene, exhibited characteristics suggestive of specificity to a persistent, non-HIV pathogen. The presence of both infected and uninfected cells in this clone at the earliest sampling (<4 months post-HIV acquisition) indicates clonal expansion prior to HIV infection. Persisting over >12 years, it peaked at frequencies above 1 per 1,000 total CD4^+^ T-cells (**Fig. 1d**). These clues prompted us to test for responsiveness to CMV (HHV-5) and EBV (HHV-4), and we observed specificity for the latter (**Fig. 1e**). Moreover, ARC ZNF721 robustly degranulated granzyme B in response to EBV antigenic stimulation indicating a cytotoxic CD4^+^ T-cell functional profile (**Fig. 1e**). These observations illustrate how purified ARCs can provide insights into the immunological context of reservoir persistence.

When co-cultured with activated CD4^+^ T-cells, each of the intact-provirus ARCs tested (n=4) seeded HIV replication, confirming their capacity to drive virologic rebound and validating their relevance as a model of HIV persistence (**Fig. 1F, Extended data Fig. 1)**. For ARC Chr22, this aligns with previous detection of the corresponding virus in a viral outgrowth assay from *ex vivo* CD4^+^ T-cells^17^.

### Transcriptomic and immunophenotypic features of ARCs

We performed multimodal single-cell sequencing (5’ CITE-seq)^41,42^ on ARCs and autologous uninfected clones that had been isolated and cultured in parallel. Initially, we analyzed synchronously isolated ARC Chr16, dARC COG5, and two uninfected clones from donor OM5334 at 3 weeks of culture, with an additional analysis of ARC Chr16 sampled after 5 months of culture to assess the stability of clonal characteristics. Clustering was primarily driven by clonal identity, and ARC Chr16 exhibited remarkably stable characteristics across timepoints (**Fig. 2a, b and Extended data Fig. 2a**). ARC Chr16 expressed high levels of *FOXP3* transcripts and elevated surface CD25/CD39, consistent with a T_reg_ phenotype. dARC COG5 exhibited high *IFNG* transcript and surface CXCR3 levels, indicating a T_H_1 profile, whereas one uninfected clone expressed *IL-4*/*IL-5* transcripts, indicative of a T_H_2 phenotype and a second had a T_H_1 profile (**Fig 2b,c and Extended data Fig. 2a**). Notably, both dARC COG5 and ARC Chr16, along with the T_H_1 uninfected clone, shared elevated perforin transcripts compared to the T_H_2 uninfected clone (**Fig 2b**). ARC Chr16 demonstrated two pronounced subclusters, with one marked by *MKI67* expression, indicating active proliferation (**Fig 2c,d**). Interestingly, both of these subclusters were present at both ARC sampling timepoints and at very similar proportions, and the UMAPs were nearly superimposable (**Extended data Fig. 2b**). Thus, even given some intra-timepoint diversity, the multiomics profile of this ARC remained remarkably stable. Pathways enriched in dARC COG5 included interferon signaling and KEAP1-NERF2 oxidative stress responses (**Fig. 2d, Extended data Fig. 10**). These results highlight that ARCs exhibit diverse and distinctive foundational characteristics that stably persist through long-term *in vitro* propagation.

**Figure 2.**
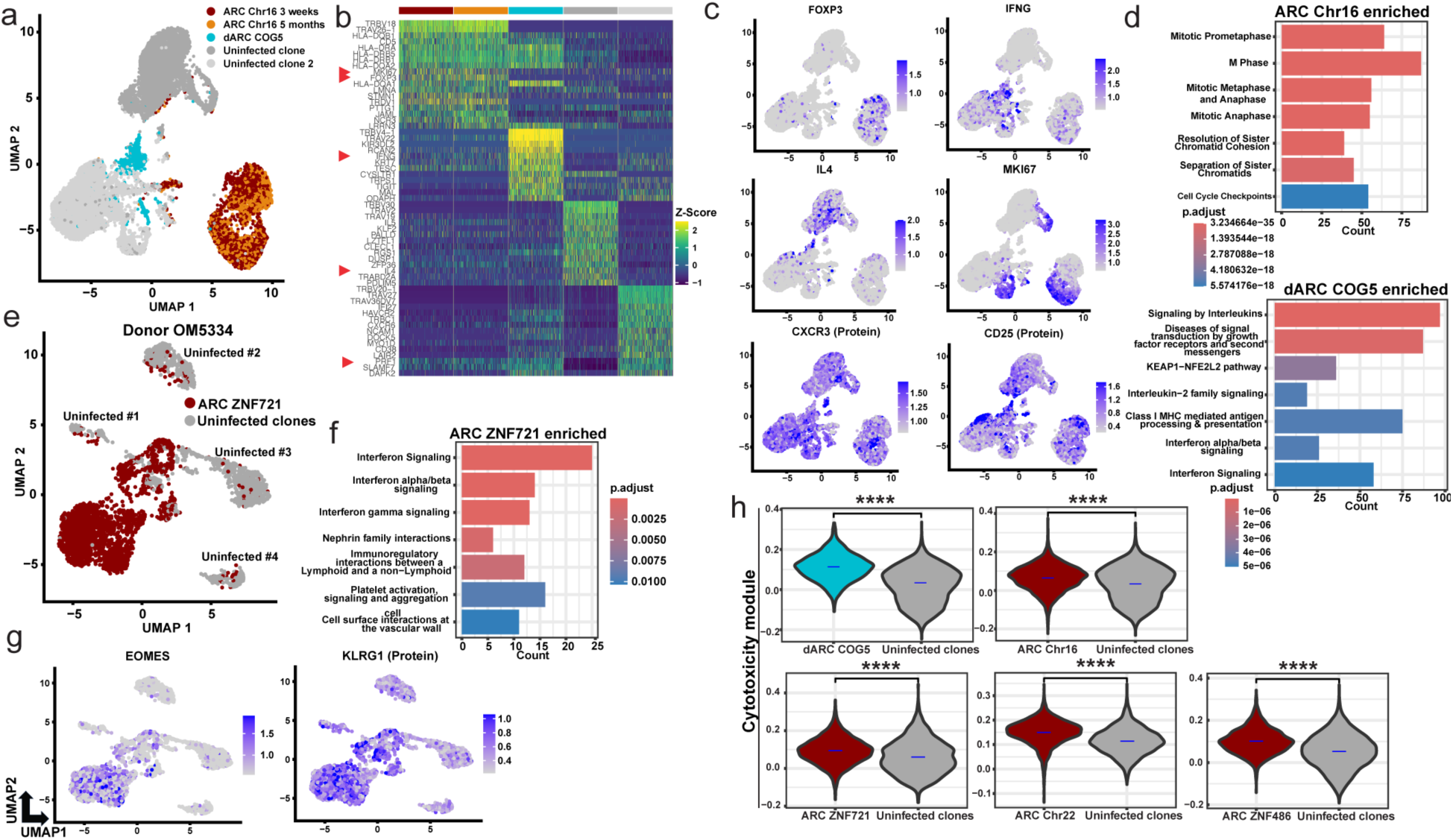
Multimodal profiling of immunophenotypic features of ARCs. **(a**) UMAP clustering of weighted nearest neighbor (WNN)-based integrated gene expression and surface protein data from OM5334-derived uninfected, dARC COG5, and ARC Chr16 clones. ARC Chr16 is compared across an early (3 weeks post-isolation) and a late (5 months) timepoints. **(b)** Heatmap of top 15 differentially expressed genes (DEGs) from clones in (a). Red arrows indicate T_H_ lineage or cytotoxic cell-related genes. **(c)** Expression of *FOXP3, IFNG*, *IL4* and *MKI67* transcripts and CXCR3 and CD25 surface proteins from clones in (a). **(d)** Top 7 enriched REACTOME pathways (merged time points) for ARC Chr16 (top) and dARC COG5 (bottom) compared to uninfected clones. **(e)** UMAP clustering of integrated gene expression and protein data from ARC ZNF721 and 3 synchronous uninfected clones, colored by infection status. **(f)** Top 7 enriched REACTOME pathways in ARC ZNF721 compared to uninfected clones. (**g**) Expression of *EOMES* and surface KLRG1 protein from clones in (**e**). (**h**) Cytotoxicity module scoring (CD8 Cytotoxic gene-set from Szabo *et al*., 2019^43^) of ARCs versus synchronous uninfected clones. Numbers of uninfected clones were: COG5 – 2, Chr16 - 2, ZNF721 – 4, Chr22 – 2, ZNF486 – 3. P-values shown are from a Wilcoxon test, **** denotes P < 0.0001.

We next expanded the analysis to ARC ZNF721 from donor OM5334 comparing to 4 synchronously isolated and cultured uninfected clones. ARC ZNF721 expressed *IFNG* transcripts, elevated cytotoxicity-associated genes (*EOMES, GZMK*), interferon signaling genes, and interferon-stimulated genes (*APOBEC3G, TRIM22*), along with surface KLRG1 protein, indicating a cytotoxic T_H_1 profile (**Fig. 2e-g, Extended data Fig. 2c-e**). The transcriptional profiles of cell lines enriched for the reservoir clones Chr22 (donor 5104) and ZNF486 (donor 603), showed these as having elevated cytotoxic transcripts (*GZMK, GZMA*), clustering near autologous *ex vivo* cytotoxic CD4^+^ T-cells^36^. Analysis of surface protein profiles from these ARCs by CITE-seq revealed overexpression of cytotoxicity/NK-cell associated CD56 surface protein in ARC Chr22 as well as overexpression of MHC-II (HLA-DR) and CD86 on both ARC Chr22 and ZNF486 compared to synchronous uninfected clones, indicative of increased IFN-γ signaling and a T_H_1 profile (**Extended data Fig. 2f,g**). Functional T_H_ phenotypes from 4 ARCs were confirmed by cytokine analysis, verifying T_H_1 profiles for dARC COG5, ARC ZNF486 and ARC ZNF721 (98.3%, 97.6% and 99.2% IFN-γ high producing-cells respectively) and a T_reg_ subtype for ARC Chr16 (45.7% of cells FoxP3^high^) (**Extended data Fig. 2h,i**). A module-scoring approach using a published cytotoxic T-cell gene-set confirmed significant overexpression of the cytotoxic module in each ARC compared to uninfected clones^43^ (**Fig 2h**). This finding aligns with recent reports identifying cytotoxic profiles as a hallmark of *ex vivo* latent and active HIV reservoirs^36,44^. Collectively, our analyses reveal diverse cellular programs across ARCs, predominately T_H_1-like but also including a distinct T_reg_ ARC, yet all are unified by enhanced cytotoxic features relative to uninfected clones.

### ARCs proliferate while appearing predominately latent

HIV transcription is facilitated by host factors that are induced in activated CD4^+^ T-cells, including NF-κB, NFAT, SP1, AP-1, BARF, IRF4^45,46^. As such, proviral latency is closely associated with resting CD4^+^ T-cells^47^. Yet, recent *ex vivo* studies infer that cells harboring intact proviruses can proliferate without viral reactivation, generating progeny cells that produce virus upon subsequent stimulation^30,48,49^. This uncoupling of proliferation from latency reversal is thought to enable HIV persistence, but scarcity of cells harboring intact HIV proviruses makes this exceptionally challenging to study. Quantification of the frequencies of cells actively transcribing HIV *ex vivo* has been achieved for three large clones with intact proviruses, yielding estimates of only 1.2-8.8%^50^. Crucially, no prior study has directly linked quantitative measures of viral expression with the activation and proliferation status of a reservoir clone.

We leveraged ARCs to address this important gap, first assessing HIV expression in the presence or absence of potent T-cell stimulation. Mining our CITEseq data, we detected HIV transcripts in the following frequencies of cells: ARC Chr16 – 5.7%, dARC COG5 – 3.1%, ARC ZNF721 –1.2% (**Fig. 3a**). We then focused on HIV-Gag protein expression by flow cytometry both for its single-cell resolution and its relevance as a prominent target of CTLs, adopting a dual anti-Gag staining strategy to enhance sensitivity and specificity (**Fig. 3b**)^51^. Across the panel of ARCs, basal HIV-Gag expression ranged from 0.0% for ARC ZNF486 to - 3.4% for ARC Chr22 (**Fig. 3b,c**), aligning with the relatively low and higher expression of these reservoir clones in enriched T-cell lines^35^. Frequencies of HIV-Gag-expressing cells were also generally in line with frequencies of cells with detectable HIV transcripts by CITEseq (though for comparisons of individual ARCs, note that sampling timepoints are not matched). In parallel, we assessed HIV-Gag expression in ARCs stimulated with anti-CD3/CD28, which mimics conventional TCR engagement, or with a combination of phorbol 12-myristate 13-acetate and ionomycin (PMA/I). Each of these strongly activate transcription factors NF-κB, NFAT, and AP-1 and are generally considered to achieve ‘maximal’ HIV reactivation^9,52,53^. While these stimuli modestly increased HIV-Gag expression in some ARCs (e.g. from 0.4% baseline to 0.9% anti-CD3/CD28 and 1.0% PMA/ionomycin for ARC ZNF721; and from 0.0% baseline to 0.05% for each stimuli for ARC ZNF486), in ARC Chr22 they decreased frequencies of HIV-Gag-expressing cells (**Fig. 3c**). Across all ARCs, most proviruses remained remarkably latent, despite potent T-cell activation (as evidenced by degranulation and release of granzymes, **Extended data Fig. 4**).

**Fig. 3.**
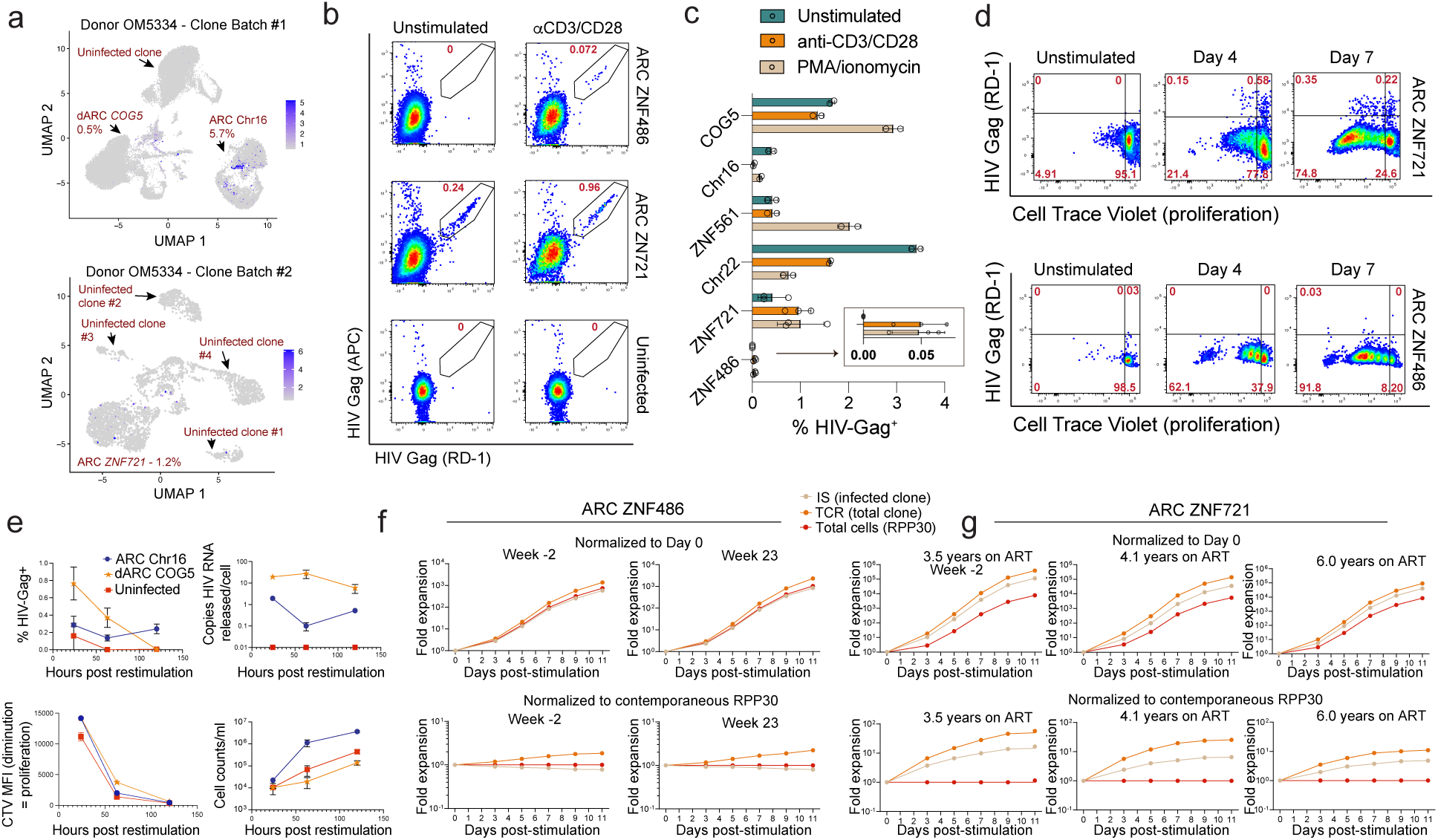
HIV expression and proliferation dynamics of ARCs in response to stimulation. **(a)** UMAP projections of integrated RNA and protein expression illustrating single-cell HIV RNA expression in ARC populations from donor OM5334. Percentages denote the proportion of cells expressing >2 log-normalized HIV RNA expression. (**b**) Flow cytometry quantification of HIV-Gag in ARCs stimulated with anti-CD3/CD28 Dynabeads, versus unstimulated. (**c**) Summary data of Gag expression, extended to PMA/ionomycin stimulation (mean±SD). (**d**) ARC proliferation following stimulation with anti-CD3/CD28 and feeder cells as assessed by CellTrace Violet (CTV) dilution on days 4 and 7; compared to unstimulated ARCs ZNF721 and 486. (**e**) Kinetics of HIV-Gag expression, viral RNA release, cellular proliferation (inverse CTV mean fluorescence intensity, MFI), and total cell counts over time following restimulation (ARC Chr16, defective ARC COG5, and uninfected control clone shown). (**e–f**) Proliferation of donor-derived T-cells stimulated *ex vivo*. Integration site (IS)-specific PCR quantifies the infected clone by detecting the proviral integration site. TCR measures the rearranged T-cell receptor of the T-cell clone (detects infected and uninfected subpopulations). Data show fold-expansion normalized to baseline (day 0, upper panels) and to contemporaneous RPP30 quantification (lower panels), for ARC ZNF486 (**e**) and ARC ZNF721 (**f**) at indicated sampling timepoints on ART.

To examine proliferation alongside HIV expression, we labeled ARCs with CellTrace Violet (CTV) and activated them by TCR stimulation. In ARC ZNF721, we observed robust proliferation that was remarkably uncoupled from latency reversal, with only 0.73% of cells expressing HIV-Gag on day 4 and 0.57% on day 7 (**Fig. 3d**). We extended this analysis to a trio of CD4^+^ T-cells clones isolated contemporaneously from donor OM5334: i) intact ARC Chr16, ii) defective ARC COG5, and iii) an uninfected clone, and longitudinally probed HIV expression, proliferation rate, and cell accumulation following a single round of restimulation with irradiated feeder cells and anti-CD3/CD28. ARC Chr16 displayed the highest increase in total cell counts reflective of its proliferative profile in our transcriptomics studies (**Fig.2b-d**), but as in previous experiments, only a fraction of the intact ARC (<0.5%) expressed HIV-Gag. This frequency was stable from 24-120 hours post restimulation and mirrored by the release of HIV RNA into supernatants. The defective ARC showed highest levels of HIV-Gag expression 24 hours post-restimulation when then declined steadily. While CTV diminution assays indicate similar levels of proliferation across the trio, ARC Chr16 harboring the intact virus accumulated to substantially greater cell numbers than either the uninfected or defective provirus clones (**Fig. 3e**), aligning with the proliferative signature evident in the multiomics characterization (**Fig. 2d**).

We further compared the proliferation of ARCs ZNF486 and ZNF721 (donors 603 and 5334, respectively) to autologous uninfected CD4^+^ T-cells clones that had been isolated and cultured in parallel. Across multiple wells from the subcloning plates, both ARCs ZNF486 and ZNF721 proliferated at rates comparable to uninfected clones (**Fig. 3d, Extended data Fig. 3**). However, these experiments are limited by the relatively few uninfected clone comparators and by the possibility that *in vitro* culture altered the expansion properties of ARCs. We therefore measured the frequencies of ARCs in total CD4^+^ T-cell populations with our integration site- and TCR Vβ-specific ddPCR assays to assess their ability to expand *ex vivo* after robust T-cell stimulation (anti-CD3/CD28). Relative to total CD4^+^ T cells, ARC ZNF486 expanded at a similar rate (**Fig. 3f**), consistent with our findings that it has a comparable proliferative capacity to that of the average autologous uninfected CD4^+^ T-cell clone. In contrast, ARC ZNF721 markedly outcompeted total CD4^+^ T-cells in the same culture, increasing by 8.7-fold in relative frequency by day 11 averaged across *in vivo* time points (**Fig. 3g**). Notably, cells with an ARC TCR clonotype (including uninfected cells of that TCR clonotype) expanded to greater numbers than cells with the paired HIV integration site (2.6-fold and 3.3-fold relative increase for ARC ZNF486 and ARC ZNF721 averaged across *in vivo* time points), suggesting a modest impact of viral cytopathic effects. Collectively, these findings demonstrate that ARCs possess robust proliferative capacities comparable to or exceeding those of autologous uninfected CD4^+^ T-cell clones, further underscoring proliferative capacity as a feature contributing to the persistence and expansion of HIV reservoirs.

### Vulnerability of Proliferating ARCs to Elimination by CTL

The prevailing HIV reservoir elimination strategy, termed ‘shock and kill’, involves pairing latency-reversal, to induce viral antigen expression, with immune effectors that eliminate reactivated cells^54^. Our observation that only a small fraction of ARC cells express HIV-Gag at any given time, even following TCR and mitogenic stimulation, would seemingly undermine the potential for their elimination by CTLs. However, we hypothesized that these static snapshots might mask dynamic antigen expression patterns that could expose larger fractions of ARCs to CTL-mediated clearance. To first assess the plausibility of this idea, we asked whether HIV-Gag expression in ARC Chr16 was a stable or transient feature of individual cells. Cells were either enriched or depleted for those expressing HIV by leveraging CD4-downregulation on HIV-expressing cells for magnetic cell-sorting. The CD4-depleted fraction exhibited the expected enrichment for HIV-expressing cells (16.8% HIV-Gag^+^) whereas the CD4-enriched fraction was depleted of HIV-expressing cells (0.09% HIV-Gag^+^) (**Fig. 4a**). Five days later, HIVGag expression levels between these two populations had converged appreciably to 0.33% in the previously depleted fraction and 4.29% in the previously enriched fraction. By day 10, these reached 0.37% and 1.10%, respectively. These results are consistent with dynamic processes of reactivation from latency and of either death or re-entry into latency of HIV expressing cells.

**Figure 4.**
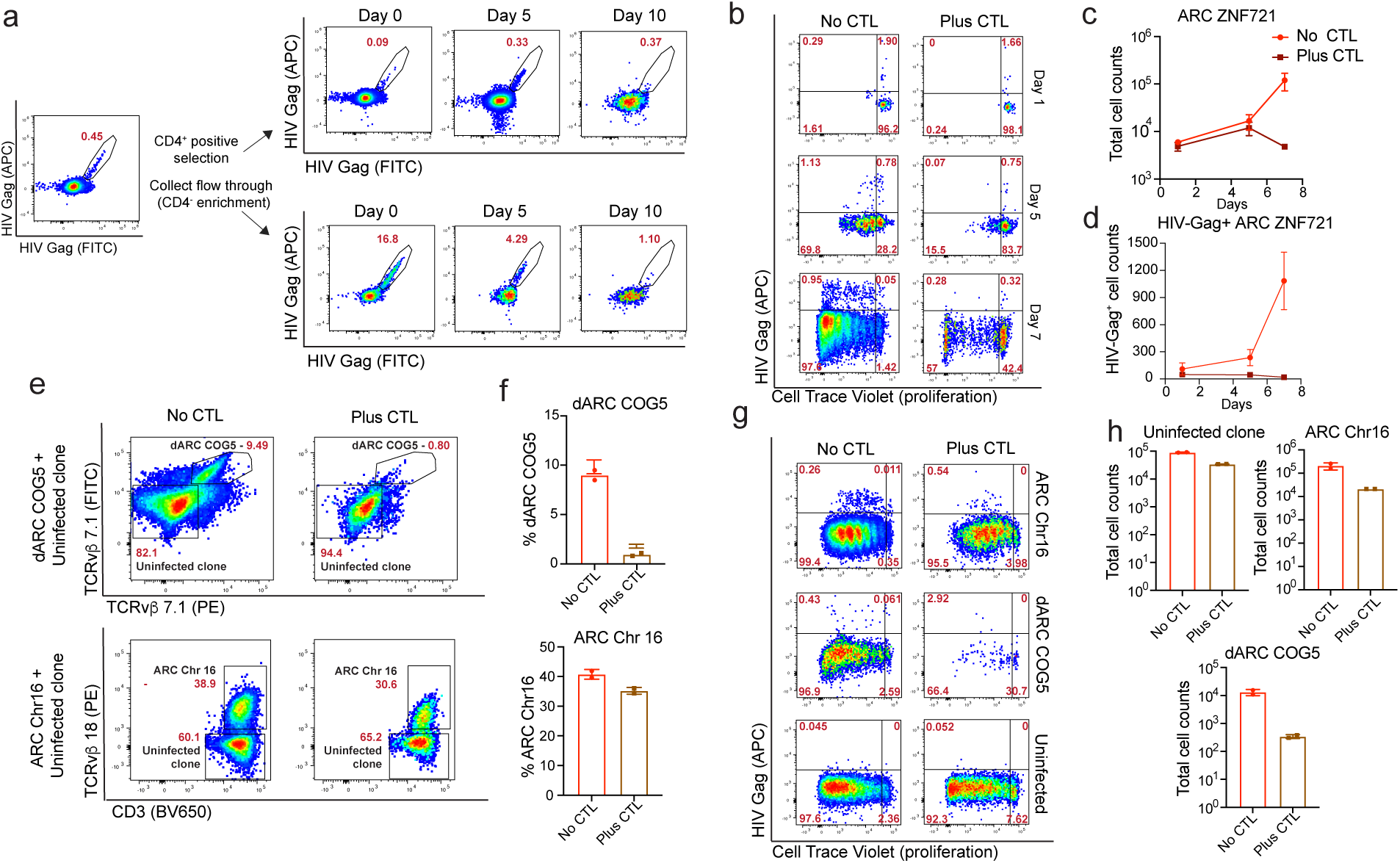
Sustained CTL pressure reveals antigenic vulnerability and intrinsic resistance of HIV reservoir clones. **(a)** Assessment of dynamic HIV-Gag expression in ARC Chr16 following initial depletion (CD4-positive selection) or enrichment (flowthrough) of HIV-expressing cells. HIV-Gag expression was monitored longitudinally at days 0, 5, and 10 post-selection by flow cytometry. (**b**) Representative slowcure assay flow cytometry plots showing HIV-Gag expression and proliferation (CellTrace Violet dilution) of ARC ZNF721 ± autologous HIV-specific CTLs (effector:target ratio 1:2). **(c-d)** Total (**c**) and HIV-Gag^+^ (**d**) ARC ZNF721 counts over 7-day co-culture ±CTLs. **(e,f**) Results from 7-day slowcure of mixed populations of ARC/dARCs with uninfected clones. dARC-COG5 is identified by dual staining with TCRvβ 7.1 antibodies versus with TCRvβ 18 for ARC Chr16. (**g,h**) Proliferation and HIV-Gag (g) and quantitative data (h) for slowcure assays. dARC COG5 is from the mixed culture in e, gated on TCRvβ 7.1 cells. The ARC Chr16 and uninfected clones are from monocultures ±CTLs setup in parallel.

In light of this evidence for dynamic HIV expression, we next evaluated the impact of both immediate and sustained CTL pressure by performing ARC proliferation assays in the presence or absence of autologous HIV-specific CTL clones (‘slowcure’ assay). We first applied this assay to ARC ZNF721 from donor OM5334. As expected, overnight culture with an HIV-Pol-specific CTL clone caused only a modest reduction in the small fraction of HIV-Gag^+^ cells without substantially affecting the total ARC population (**Fig. 4b,c**). By day 5 of co-culture, we began to observe significant reductions in both HIV-Gag^+^ cells and total ARC numbers, particularly those that had undergone proliferation. By day 7, CTLs had reduced surviving ARCs by >90%, despite fewer than 1% of cells expressing Gag at each sampling snapshot. A similar pattern was observed in ARC Chr22 from donor 5104, though less proliferation was observed (**Extended data Fig. 5**).

In extending these results to a second intact ARC (Chr16) from donor OM5334, we observed a strikingly different result, with only modest reduction in either the frequencies of HIV-Gag^+^ cells or total ARC cells at days 1, 4, and 7 of co-culture with the same HIV-Pol-specific CTL clone (**Extended data Fig. 6**). The epitope targeted by this CTL clone (NETPGIRY, HLA-B*18-restricted) was confirmed to be intact in the ARC Chr16 virus. Results were suggestive of a modest CTL impact: i) whereas the CTL enhanced expansion of an uninfected CD4^+^ T-cell clone, there was a trend towards lower ARC numbers at day 7, and ii) MHC-I blocking experiments suggested limited reductions in frequencies of HIV-Gag^+^ cells. However, in contrast to the result with ARC ZNF721, the overall outcome was robust expansion of ARC Chr16 despite the presence of CTLs (**Extended data Fig. 6**).

To further investigate this finding, we conducted slowcure assays comparing ARC Chr16 and its autologous defective counterpart dARC COG5 (5’-L defective). We included internal controls of autologous uninfected CD4^+^ T-cell clones in the same co-cultures as the ARC/dARC. For dARC COG5, we used a partially subcloned (ARC precursor) sample containing a mixture of the defective clone and an uninfected clone, ensuring identical cell culture histories. For ARC Chr16, we simulated this by spiking in an autologous uninfected clone that had been isolated and cultured in parallel. The clones were distinguished in each culture by their distinct TCR Vβ segments (Vβ7.1 for dARC COG5 and Vβ18 for ARC Chr16). Over the 7-day CTL co-culture, dARC COG5 was potently and specifically eliminated (**Fig. 4e,f**). In contrast, ARC Chr16 showed minimal selective elimination compared to its matched uninfected control (**Fig. 4e,f**). Monocultures of ARC Chr16 and uninfected clones, analyzed alongside TCR Vβ7.1-gated dARC COG5, further corroborated the relative resistance of ARC Chr16 to CTL-mediated killing (**Fig. 4g,h**). Thus, despite transient HIV-Gag expression, ARC ZNF721 and dARC COG5 exhibited pronounced vulnerability to CTL elimination, whereas ARC Chr16 was minimally impacted. The persistence of an HIV-Gag^+^ suggests that latency may not be the only barrier to CTL killing. population in ARC Chr16 under sustained CTL pressure suggests that latency alone does not fully explain resistance to CTL-mediated killing.

### ARC Resistance to CTL Killing and Reversal by Hypoxia Induction

To determine whether the limited susceptibility of ARC Chr16 to CTL was due to viral or cellular factors, we tested whether the Pol-specific CTL clone could kill bulk autologous CD4^+^ T-cells infected with virus derived from ARC Chr16. By differentially labeling these newly *in vitro* infected cells, dARC COG5, and ARC Chr16 we assessed susceptibility to CTL of HIV-Gag^+^ cells in these three types of autologous CD4^+^ T-cells to elimination in competitive overnight killing assays (**Fig. 5a)**. We observed potent killing of HIV-Gag^+^ dARC COG5 cells, contrasted by a lack of any reduction in HIV-Gag^+^ ARC Chr16 cells, aligning with their respective results in the slowcure assays (**Fig. 5b, Extended data Fig. 7**). The majority (>75%) of bulk CD4^+^ T-cells that had been newly infected with virus from ARC Chr16 were effectively killed by CTL, indicating that the lack of elimination of HIV-Gag^+^ ARC Chr16 was attributable to cellular properties of this reservoir clone rather than to the virus it harbors. Of note, is the survival of 25% of the newly infected HIV-Gag^+^ bulk CD4^+^ T-cells, both at 1:10 and excessive 1:2 effector:target ratio (**Fig. 5b**). This is typical of a phenomenon we have recently described, resulting from a subset of CD4^+^ T-cells that are resistant to elimination by CTL^23,24,55,56^. Thus, ARC Chr16 exhibited a resistance to CTL that was associated with cellular rather than viral properties.

**Figure 5.**
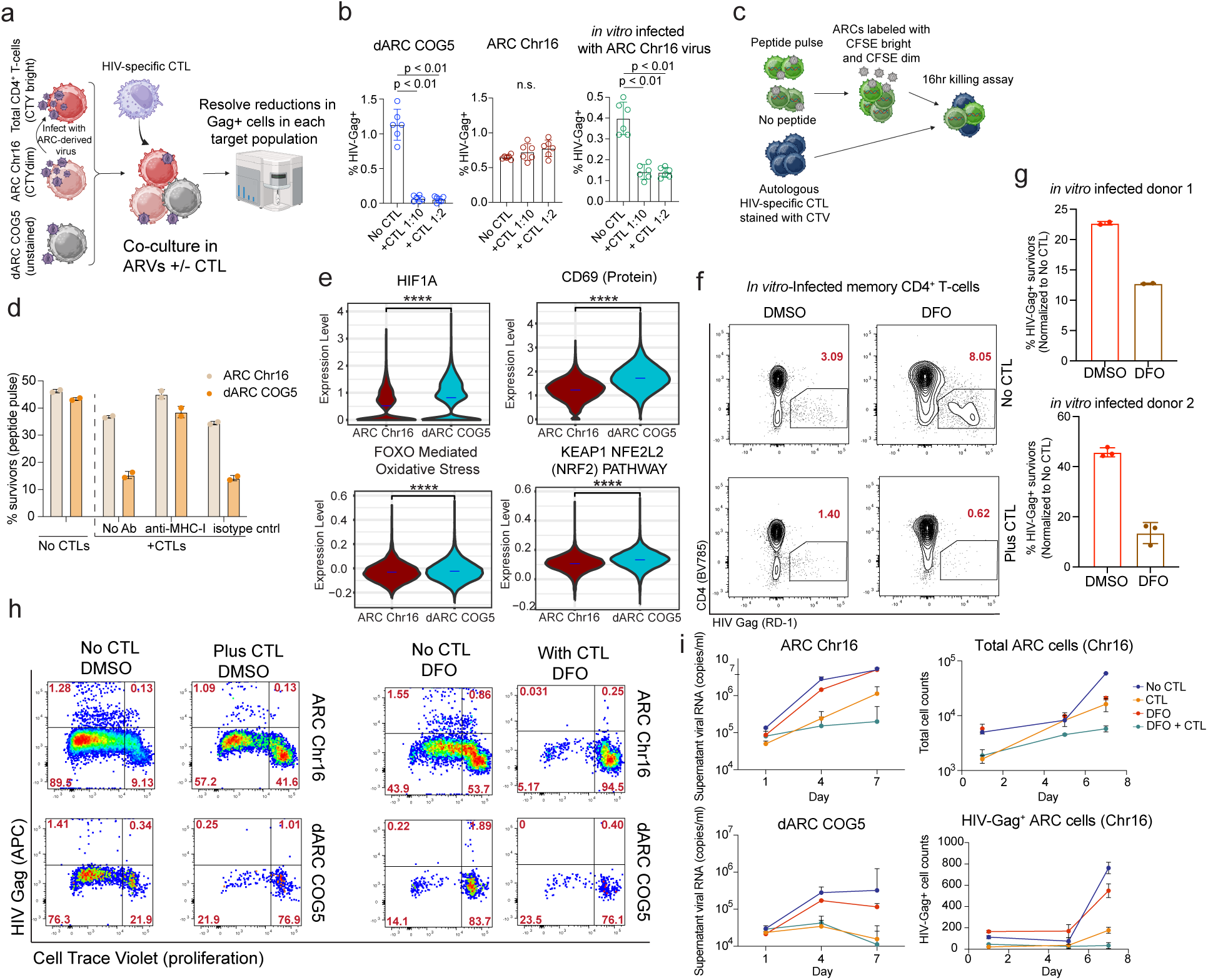
Cell-intrinsic resistance of a reservoir clone to elimination by CTL. **(a,b)** Schematic (a) and quantitative data (b) from competitive overnight killing assays measuring CTL susceptibility of HIV-Gag^+^ cells in ARC Chr16, dARC COG5, and bulk CD4^+^ T-cells infected with ARC Chr16-derived virus. Statistical analysis: Kruskal-Wallis test, Dunn’s multiple-comparison test (two-sided). (**c,d**) Schematic (c) and quantification (d) of CTL killing of peptide-pulsed/unpulsed ARC Chr16 and dARC COG5 labeled with differential CFSE intensities ±MHC-I blocking antibody. (**e**) Violin plots showing expression of *HIF1a*, surface CD69 protein, and modules scores of Reactome “FOXO mediated transcription of oxidative stress metabolic and neuronal genes” and “KEAP1 NFE2L2 pathway” gene-sets on ARC Chr16 and dARC COG5. Blue dash denotes mean expression. P-values shown are from a Wilcoxon test (two-sided), **** P < 0.0001. (**f,g**) Overnight killing assays testing the effects of deferoxamine (DFO, 72 hour treatment) on memory CD4^+^ T-cells from 2 donors *in vitro*-infected with HIV-JRSCF and co-cultured with/without Gag-TW10 specific CTL clones. (**f**) Shows representative flow cytometry plots after the killing assay from one donor, summarized by donor in (g). **(h-i)** Slowcure assay evaluating DFO on viral RNA release, total cell counts, and HIV-Gag^+^ ARC cells (Chr16, dARC COG5 ±CTL); representative flow cytometry at day 7 (**h**), quantitative data (**i**).

We asked whether the cell-intrinsic ability of ARC Chr16 to survive CTL attack were tied to a deficiency in HIV antigen processing/presentation. For both ARC Chr16 and dARC COG5, cells were divided into two aliquots and either pulsed or not with the HIV-Pol peptide recognized by the CTL clone. These were labeled with different concentrations of a CellTrace dye to differentiate based on fluorochrome brightness, mixed, and then cultured with or without the CTL clone. Whereas dARC COG5 was efficiently eliminated by CTLs in a peptide and MHC-I-dependent manner, ARC Chr16 was only marginally impacted, indicating resistance to death downstream of CTL recognition (**Fig. 5c,d**). Relative susceptibilities of other ARCs to CTL are given in **Extended Data Fig.8a**).

In studying populations of HIV-infected CD4^+^ T-cells that survive CTL coculture, we have thus far reported three mechanisms of resistance: i) overexpression of the prosurvival factor BCL-2^23^, ii) overexpression of EZH2^55^, resulting in suboptimal surface MHC-I, iii) metabolic quiescence leading to dampened levels of hypoxia and oxidative stress^56^. Staining for BCL-2 had been included in the competitive killing assay in **Fig. 5a,b**, and while both the dARC and the ARC exhibited high BCL-2 expression relative to bulk CD4^+^ T-cells, levels were highest on the former – arguing against a major role in the current observation (**Extended data Fig. 7)**. For HIV-Gag^+^ cells, MHC-I expression was downregulated on both ARC Chr16 and newly-infected targets), but not the dARC – potentially indicating that the defect impacts HIV-Nef expression. While this may contribute to the particular sensitivity of the dARC to CTL, it cannot explain the discordance between ARC Chr16 and newly infected targets – and survivors from the latter did not show selection for MHC-I-low cells; nor does it fit with the insensitivity of peptide-pulsed ARC Chr16 to CTL (HIV-Gag^-^ cells from ARC Chr16 show normal levels of MHC-I).

To identify other mechanisms by which ARC Chr16 may resist CTL, we cross-referenced our CITE-seq analysis of ARC Chr16 and dARC COG5 with our recent multiomic profiling of cells that survive CTL killing in an *in vitro* infection system^56^. In this model system, relatively quiescent (CD69^low^) cells with reduced expression of the hypoxia-promoting *HIF1A* and lower oxidative stress responses genes preferentially survived CTL co-culture, likely due to the role of reactive oxygen species (ROS) in death downstream of a granzyme/perforin hit^57,58^. Aligning with this, ARC Chr16 showed significantly lower levels of *HIF1A*, surface CD69 expression as well oxidative stress response modules compared to the CTL-vulnerable dARC COG5 (**Fig. 5e, Extended data Fig. 10**). The FDA-approved therapeutic deferoxamine (DFO) counteracts these resistance-associated features by inducing hypoxic stress and ROS^56^. DFO treatment of *in vitro* HIV-infected CD4^+^ T-cells enhanced CTL killing in an overnight culture (**Fig. 5f,g**), cutting into the resistant fraction that we consistently observe (**Fig. 5b,d)**. We assessed whether the induction of hypoxic stress on ARC Chr16 with DFO during a slowcure assay would facilitate killing by CTL. As previously, without DFO CTL modestly impacted accumulation of ARC Chr16, allowing it to expand 3.6-fold from baseline (versus 11.8-fold without CTL). With the addition of DFO to the CTL co-cultures, proliferating ARCs were largely eliminated, allowing for only a 1.1-fold increase (**Fig. 5h,i**). This was mirrored by viral RNA in supernatants, which increased to 40-fold by day 7 in the absence of CTL, 7.6-fold in the presence of CTL alone, and 1.8-fold with the combination of CTL and DFO. Thus, knowledge of both the characteristics of the ARC Chr16 and of CTL resistance mechanisms in HIV-infected CD4^+^ T-cells facilitated a targeted elimination strategy.

## Discussion

By isolating ARCs from *ex vivo* from CD4^+^ T-cells, we are continually expanding a resource that enables unprecedented immunologic, virologic, and functional characterization of persistent HIV reservoirs. The ARC described here are as diverse as T-cells themselves, yet each skew towards a cytotoxic phenotype, aligning with recent *ex vivo* observations^36,44^. We further provide the first functional confirmation of this profile, showing high basal levels of granzymes in ARCs and their release upon stimulation. These ARCs overtly manifest features underpinning their notorious resistance to HIV cure strategies. They proliferate robustly with only transient viral expression – sufficient to seed new infections – but show minimal sensitivity to the ‘shock’ phase of the conventional ‘shock and kill’ approach, which envisions broad pharmacological induction of viral antigens. Nonetheless, during proliferation, three of four ARCs tested demonstrated vulnerability to CTL-mediated attack. This suggests a model in which the interception of spontaneous and transient HIV expression may be sufficient to allow for gradual reservoir elimination, given sufficiently potent CTL responses. Indeed, the pronounced erosion of rebound-competent reservoirs observed in ‘elite controllers’ may reflect this mechanism, given their hallmark robust CTL responses^19^. However, despite potent CTL interception of proliferating ARCs, we consistently observed that some cells persisted through multiple rounds of proliferation. We also consistently observed subsets of non-proliferating survivors. Further study is needed to understand the basis of both of these modalities of survival, and whether complete elimination can be achieved through either improved latency reversal or CTL sensitization strategies. Additionally, evaluating CTL efficacy against ARCs independent of TCR stimulation, for example using latency reversal with Tat-containing lipid nanoparticles, will be critical^59,60^.

To what extent do our isolated ARCs represent the broader reservoir? Our methodological approach – particularly limiting dilution – may preferentially isolate reservoir clones adept at proliferating while expressing HIV-Gag protein. These clones may represent amongst the most refractory reservoir subsets that retain the ability to re-seed replication. Indeed, we observed integration site characteristics in ARCs (in ZNF genes and heterochromatic regions) converging with those of reservoir clones persisting long-term under immune pressure *in vivo* and notably, the remarkable 12-year *in vivo* stability of the OM5334 ARCs^19^. Alternative cloning strategies (e.g. screening by HIV PCR rather than p24 ELISA to detect deeply latent clones) could unveil additional ARC subtypes.

We directly demonstrate cell-intrinsic CTL resistance in a reservoir-harboring clone - in this case a T_reg_. Such resistance was initially hypothesized based on the minimal impact on the reservoir with combinations of potent LRAs and CTLs *ex vivo* studies^24^. Proposed mechanisms have included the granzyme B inhibitor SERPINB9, the inhibitory ligands PVR^22,44^, and overexpression of the anti-apoptotic protein BCL-2 – inhibitors of which facilitate reservoir reductions *ex vivo* and in non-human primates^23,61^. Our direct demonstration of intrinsic CTL resistance provides validation and impetus to accelerate the identification of diverse resistance pathways within HIV reservoirs. Building upon the precision approach used against ARC Chr16, further study is needed to determine the degree to which DFO or other agents (alone in combination) can broadly sensitize HIV reservoirs clones to elimination, opening avenues for HIV cure strategies beyond traditional latency reversal approaches

Given that cell-intrinsic resistance to CTL limits cancer immunotherapies, our findings pave the way for cross-disciplinary insights. Previously reported mechanisms of CTL resistance in HIV - including BCL-2 and EZH2 overexpression - are already recognized therapeutic targets in leukemia and lymphoma^62,63^, while DFO remains unexplored in immunotherapy contexts. Ultimately, our ARC isolation platform has enabled us not only to directly demonstrate critical therapeutic vulnerabilities and resistance mechanisms within HIV reservoirs but also provides a framework for probing fundamental aspects of human T-cell biology – particularly those governing proliferation and immune resistance – with implications for and beyond HIV persistence.

## Acknowledgments

We thank all study participants who devoted time to our research, as well as the clinical research team involved in the study. This work would not have been possible without regents provided by the AIDS and Cancer Virus Program, Leidos Biomedical Research, Inc., Frederick National Laboratory for Cancer Research, supported with federal funds from the National Cancer Institute, National Institutes of Health, under contract HHSN261200800001E. We acknowledge the Biological Resources Branch Preclinical Repository, National Cancer Institute, for providing IL-2 and IL-15. We thank Adam R. Ward for his review of the manuscript and useful discussion. This work was supported by the following NIH/NIAID grants: R01MH130197, R01AI170239, R37AI181626, R01AI176601, R01AI176943, R01AI170245, R01AI165031, UM1AI164562, UM1AI164565, R01AI150412, R21AI170246, R01AI147845, U01AI145921, R01AI167691, R21AI172554 and 1F31AI191953-01. UM1 AI16452 and UM1AI164565 were also supported by NIMH, NIDA, NINDS, NIDDK, and NHLBI. C.O. was supported by a Medical Scientist Training Program grant from the National Institute of General Medical Sciences of the National Institutes of Health under award number T32GM152349 to the Weill Cornell/Rockefeller/Sloan Kettering Tri-Institutional MD-PhD Program. M.C.N and P.B are HHMI Investigators. The funders had no role in study design, data collection and analysis, decision to publish, or preparation of the manuscript. Figure and figure schematics were created using GraphPad Prism, BioRender.com, and Adobe Illustrator.

## Author contributions

R.B.J, I.A.T.M.F., A.H, N.L.B. conceived the study. I.A.T.M.F., A.H, R.B.J, N.L.B, T.T.H., N.L.L, Y.R., C.B, G.Q.L. F.L. designed experiments. I.A.T.M.F., A.H, N.L.B, T.T.H., N.L.L, R.B.J, Y.R., C.B, V.K.P., S.V., E.N., P.S., A.D., Marie C., C.O. F.L. performed experiments. G.Q.L, R.B.J, I.A.T.M.F., A.H, N.L.B, T.T.H, A.D, P.B. analyzed and interpreted the data. C.B, M.C.N, M.C. contributed essential reagents or analytical tools. M.C., S.M., T.-W.C., C.K. provided clinical samples or patient data. A.H, P.Z., D.B., G.Q.L, V.K.P. performed bioinformatic analyses. R.B.J, P.B, M.C.N. supervised research. R.B.J, A.H, I.A.T.M wrote the manuscript with input from all authors. All authors reviewed, edited, and approved the final manuscript.

## Methods Section

### Human Samples

Peripheral blood or leukapheresis samples were obtained from individuals living with and without HIV. Participants living with HIV who had been suppressed on antiretroviral therapy to undetectable levels for at least 2 years, were recruited from the Maple Leaf Medical Clinic (Toronto, Canada), the Rockefeller University (New York, USA), or through a clinical protocol approved by the Institutional Review Board of the National Institutes of Health (NIH). Leukapheresis products were collected in accordance with approved protocols, and all participants provided written informed consent for subsequent testing. Samples from deidentified HIV-negative individuals were obtained from the New York Blood Center, the Gulf Coast Regional Blood Center, or Stemcell Technologies. The study was approved by the Institutional Review Boards of the University of Toronto, Rockefeller University, and Weill Cornell Medicine.

### Media and reagents

The following culture media were used: R10, which is RPMI 1640 (Thermo Fisher) supplemented with 10% FBS (Gibco™) and 1% penicillin/streptomycin (Gibco™), 1% HEPES buffer (Gibco™), 1% L-Glutamine (Gibco™); ‘clone media’ which is R10 supplemented with 100 Units/mL of IL-2, 1 µM Nevirapine, 1 µM Emtricitabine, 1 µM Tenofovir, 10 µM Enfuvirtide, and 10 µg/mL 10-1074 bNAb – each from the NIH AIDS reagent program; R10-50 + IL15, which is R10 supplemented with 50 Units/mL of IL-2 and 10 ng/mL of IL-15. Recombinant human IL-2 and IL-15 were obtained from National Cancer Institute.

### p24 Enzyme-linked Immunosorbent Assay (ELISA)

High-binding 96-well plates (Grenier Bio-One, Lot #655061) were coated with 100 µL per well of mouse anti–HIV p24 monoclonal antibody (AIDS and Cancer Virus Program, Frederick National Laboratory for Cancer Research) diluted in DPBS according to the manufacturer’s instructions and incubated overnight at 4°C. The next day, coating solution was removed, and wells were blocked with 200 µL of blocking buffer (PBS + 1% BSA) for 30 minutes at room temperature. Blocking buffer was then aspirated, and 100 µL of culture supernatant from each well was added along with 10 µL of lysis buffer (distilled water + 10% Triton X-100). Plates were incubated at 37°C for 1 hour.

Following incubation, wells were washed five times with 300 µL of wash buffer (PBS + 0.1% Tween-20). Rabbit anti–HIV p24 polyclonal antibody (AIDS and Cancer Virus Program, Frederick National Laboratory for Cancer Research), diluted in primary antibody diluent (RPMI-1640 supplemented with 10% FBS and 2% normal mouse serum), was added at 100 µL per well. Plates were incubated for 1 hour at 37°C, then washed again as described.

Next, 100 µL of goat anti-rabbit IgG (H+L) secondary antibody (Novus Biologicals), diluted 1:20,000 in secondary antibody diluent (RPMI-1640 supplemented with 2% normal mouse serum, 5% normal goat serum, and 0.01% Tween-20), was added per well and incubated for 1 hour at 37°C. After a final wash step, 100 µL of 1-Step™ Ultra TMB-ELISA Substrate Solution (Thermo Fisher Scientific) was added to each well and incubated at room temperature for 30 minutes in the dark. The reaction was stopped by adding 100 µL of Stop Solution for TMB Substrate (BioLegend) per well.

Plates were read at 450 nm using a SpectraMax microplate reader (Molecular Devices) with SoftMax® Pro software. Clone media was used as a negative control. Wells with absorbance value more than 1.5 times the negative controls were considered p24 positive.

### Intact Proviral DNA Assay (IPDA)

IPDA was performed as previously described^7^, on DNA from CD4^+^ T-cells populations as a routine purity check every three stimulation cycles. Genomic DNA was extracted using the DNeasy Blood and Tissue Kit (Qiagen) with the elution step using nuclease-free water. DNA concentrations were quantified using Nanodrop Spectrophotometer [Thermo Scientific]. Each ddPCR reaction was composed of 1x ddPCR Supermix for Probes (no dUTPs, BioRad), 900 nM primers (Integrated DNA Technologies), 250 nM probes (ThermoFisher Scientific), 22.5 ng of genomic DNA, and nuclease-free water. Primer and probe sequences are given in Supplementary Table 3. For each sample, the HIV reaction (HIV Ψ/HIV gag Back-up with HIV env/HIV env Back-up) and RPP30 reaction (RPP30 and RPP30 Shear) were run in duplicate. For newly isolated ARCs, two different sets of HIV reactions using different primer/probe combinations (HIV Ψ with HIV env or HIV gag Back-up with HIV env Back-up) were run in duplicate to determine the primer/probe compatibility. Droplets were prepared using an Automated Droplet Generator (BioRad), cycled at 95 °C for 10 min; 45 cycles of (94 °C for 30 sec, 59 °C for 1 min) and 98 °C for 10 min, then analyzed with QX600 Droplet Reader (BioRad) using QuantaSoft software (BioRad, version 2.1).

### Authentic reservoir clone isolation

Memory CD4^+^ T-cells (mCD4) were isolated from the PBMCs by of people with HIV (PWH) by negative selection using EasySep™ Human Memory CD4^+^ T Cell Enrichment Kit (STEMCELL Technologies), plated (10 cells/well, 96-well plates), and stimulated with irradiated allogeneic HIV-negative PBMCs (1x10^6^/mL) in clone media for 21 days. Feeder PBMCs were depleted of CD8^+^ T-cells and NK cells using EasySep™ Human CD8 Positive Selection Kit II and EasySep™ Human CD56 Positive Selection Kit II (STEMCELL Technologies) before being co-cultured with mCD4 T-cells from PWH. At the end of stimulation cycle, p24 ELISA was performed on all plates. Cells from p24 ELISA-positive wells were plated in serial 3-fold dilution (96-well plates), stimulated at the same condition as aforementioned, and subjected to p24 ELISA after 21 days. After the second round of plating and ELISA, cells from p24 ELISA-positive wells were expanded under the same stimulation condition. Clonal purity of isolated infected populations was determined using IPDA and further serial dilution subs were performed as needed.

### Near-full-length HIV sequencing (FLIP-Seq)

Genomic DNA was extracted from the clones using a QIAGEN DNeasy kit. DNA was diluted to single genome levels based on Poisson distribution statistics and ddPCR results, and was subjected to single-genome amplification using Invitrogen Platinum Taq and nested primers spanning near full-length HIV (HXB2 coordinates 638-9632). All primers were previously described^2^ and can be found in Supplementary Table 3. PCR products were visualized by agarose gel electrophoresis. All near-full-length sequences (>8,000bp) were subjected to Illumina MiSeq sequencing. Resulting short reads (142bp) were de novo assembled and aligned to HXB2 to identify large deleterious deletions, out-of-frame indels, premature/lethal stop codons, internal inversions, or packaging signal deletions, using an automated in-house pipeline written in R scripting language as previously described^2^. The presence or absence of APOBEC-3G/3F–associated hypermutations was determined using the Los Alamos HIV Sequence Database Hypermut 2.0 program^3^. Viral sequences that lacked all mutations listed above were classified as intact. Sequences were aligned to the HXB2 reference genome using Geneious Prime (version 2025.1) and manual editing where appropriate. The 5’ and 3’ ends of the sequences not covered by the near-full-length PCR were extracted from the corresponding integration site reads (HXB2 coordinates 1-681 and 9686-9719 respectively). A short 53bp section of the 3’ LTR (HXB2 coordinates 9633-9685) was inferred based on the corresponding 5’ LTR sequence. To examine phylogenetic distances between sequences, a neighbor-joining tree containing all clone proviral sequences and the HXB2 reference sequence was prepared using MEGA (version 7.0).

### Integration site sequencing

Genomic DNA was extracted from the ARCs using a QIAGEN DNeasy kit. Subsequently, 2.5 μl of gDNA sample was subjected to multiple displacement amplification (MDA) with phi29 polymerase (Qiagen REPLI-g Single Cell Kit #150345) at 30° C for 4 h. DNA produced by whole-genome amplification was digested with DraI, SspI, and HpaI. The digested DNA was purified, followed by ligation of genome-walker (GW01) adapter. Viral-host junction regions were amplified using a nested PCR strategy (see Supplementary Table 4 for primers). Resulting PCR products were subjected to Illumina MiSeq sequencing. MiSeq paired-end FASTQ files were demultiplexed; small reads (142 bp) were then aligned simultaneously to human reference genome T2T and HIV reference genome HXB2. Biocomputational identification of integration sites was performed using in house R based pipeline^1^. Briefly, chimeric reads containing both human and HIV sequences were evaluated for mapping quality based on (i) HIV coordinates mapping to the terminal nucleotides of the viral genome, (ii) absolute counts of chimeric reads, (iii) depth of sequencing coverage in the host genome adjacent to the viral integration site. The final list of integration sites and corresponding chromosomal annotations was obtained using Ensembl (v86, www.ensembl.org), the UCSC Genome Browser (www.genome.ucsc.edu) and GENCODE (v39, www.gencodegenes.org).

### Integration site- and TCR Vβ-specific ddPCR assays

We designed droplet digital PCR assays that quantify total HIV LTR, the integration site of a specific provirus, and the corresponding TCRβ CDR3 of the clone containing that provirus. The primers and probe used to amplify the R-U5 region of the HIV LTR have been described previously^4,5^. Integration site-specific and TCRβ CDR3-specific primers and probes were designed for individual reservoir clones as previously described^5,6^. Briefly, integration site-specific probes were designed to span the host-5’ LTR junction with corresponding forward and reverse primers on the host genome and the U3 region of the HIV LTR respectively. Similarly, TCRβ CDR3-specific probes were designed to span the junctions between the V-D-J regions with corresponding forward and reverse primers on the V and J regions respectively. The specificity of each primer and probe set was validated using DNA from the corresponding reservoir clone and compared to DNA from the CD4^+^ T cells of HIV-negative donors and DNA from other infected and uninfected clones. A complete list of primers and probes can be found in Supplementary Table 2. Primers and probes were used at final concentrations of 900nM and 250nM respectively. Droplet digital PCR was performed with the Bio-Rad QX600 platform using the following cycling conditions: 95°C for 10 min, 50 cycles of 95°C for 30 sec and 56°C for 1 min, 98°C for 10 min, hold at 4°C. RPP30 ddPCR was performed in parallel for cell equivalent normalization as previously described^7^. Because the HIV LTR-specific primers and probe do not distinguish between 5’ and 3’ location of the amplified R-U5 region, a correction factor of ½ is applied to LTR copy number values prior to cell equivalent normalization.

### Assessing proliferation of ARCs in *ex vivo* CD4^+^ T-cells

10^7^ total CD4^+^ T cells were plated at 10^6^ cell/mL clone media and stimulated with anti-CD3/CD28 Dynabeads at a 5:1 bead:cell ratio. 24 hours poststimulation, the cells were counted and media was added to reach a final cell density of 10^5^ cells/mL. At days 3, 5, 7, 9, and 11 poststimulation, precise total cell counts were recorded, 10^7^ cells were set aside for genomic DNA extraction and media was added to reach a final cell density of 10^5^ cells/mL. Cell viability remained at >90% for the duration of the 11-day culture. LIST ddPCR was used to quantify total HIV LTRs and the integration site and TCRβ of specific ARCs at each culture time point. Expansion dynamics of total CD4^+^ T cells was determined by relative changes in cell counts over the duration of the culture. Changes in the frequency of the ARC integration site and TCRβ to total RPP30 cell equivalents over the duration of the culture were used to calculate expansion dynamics of cells harboring the ARC integration site or TCRβ.

### ELISPOTs

To perform the IFN-γ and Granzyme B ELISPOT assays, capture antibodies (clone 1-D1K for IFN-γ, clone MT28 for Granzyme B) (Mabtech) were first diluted in PBS according to the manufacturer’s instructions and plated at 100 µL per well into multiscreen IP 96-well PVDF plates (MilliporeSigma); coated plates were subsequently incubated overnight at 4°C to allow optimal antibody binding to the membrane surface, ensuring reliable cytokine capture upon stimulation.

Following this incubation period, the plates were washed five times with PBS to remove any unbound antibody and to prepare the wells for cell seeding; subsequently, 200,000 ARCs were added per well in R10 medium containing the relevant antigens or controls. Antigens being used were CMV-pp65 peptide pool at 1 μg/mL (STEMCELL Technologies), CMV-infected cell extract, EBV-infected cell extract, and negative control cell extract (for EBV-infected cells) at 10 µL per 200,000 cells (all three from Virusys). PHA at 0.25 µg/mL served as a positive control to confirm assay responsiveness, while 0.5% DMSO was used as a negative control to determine background signal levels.

The plates were then incubated at 37°C for 17 to 20 hours in a humidified incubator, allowing T cell activation and cytokine secretion to occur; after this stimulation phase, wells were washed five times with PBS to eliminate residual cells and media, thus minimizing nonspecific staining during detection. A biotinylated detection antibody (clone 7-B6-1 for IFN-γ, clone MT8610 for Granzyme B) (Mabtech) in 100 µL PBS at the manufacturer’s recommended concentration was then added to each well and incubated at room temperature for 2 hours; after this period, plates were again washed five times to remove unbound antibody before the addition of streptavidin-ALP at 1 µg/mL in 100 µL PBS per well (Mabtech), which was incubated for 1 hour at room temperature to allow sufficient binding and amplification of the biotin signal. The plates were washed again five times and developed using AP Conjugate Substrate Kit (Bio-Rad) for 15 minutes; during this time, spot formation occurred where IFN-γ had been secreted by activated T cells and captured by the plate-bound antibody complex. Following development, plates were thoroughly washed to halt the enzymatic reaction, air-dried overnight to stabilize the spots, and subsequently analyzed using a CTL ELISPOT reader (ImmunoSpot), which quantitatively measured the number and intensity of cytokine-producing cells.

### HIV transmission assay

To determine whether virus produced by ARCs was replication competent, we co-cultured ARCs with activated CD4^+^ ‘target cells’ in the absence of ARVs and assessed infection. CD4 cells were isolated from PBMCs of HIV-negative donors via negative selection (EasySep™ Human CD4+ T Cell Isolation Kit, STEMCELL Technologies), cultured in R10-50 +IL15 (1x10^6^/mL), and activated for 72 hours with ImmunoCult™ Human CD3/CD28 T Cell Activator at 25 µL/mL (STEMCELL Technologies). ARCs were washed to remove ARVs labelled with CellTrace™ CFSE at 5 µM according to the manufacturer’s protocol (CSFE, ThermoFisher Scientific), cultured in R10-50+15 for 24 hours for additional ARV washout, and then co-cultured with target cells at ratios ranging from 1:10 – 1:100 ARC:target cells in R10-50+15. In control conditions, we cultured with a CFSE-labeled uninfected CD4+ T-cell clones that had been isolated and maintained in parallel to the ARC. Negative controls were activated uninfected CD4+ T-cells not cocultured with infected populations. At days 3 and 7, samples were stained for flow cytometry using anti-CD4 two antibodies against HIV-Gag to enhance sensitivity and specificity (see below for details). The presence of HIV-Gag^+^ populations within the CFSE-negative target cell population comprised evidence for viral transmission.

### Flow cytometry, including HIV-Gag staining

Cells were stained for viability using the Live/dead dye in MACS buffer FACS buffer (PBS + 2% FBS + 1 mM EDTA) following manufacturer’s instructions and with other surface antibodies depending on the experiment, including: CD3 OKT3 (Biolegend), CD4 OKT4 (Biolgened), CD8 SK1 (Biolegend), CD107a H4A3 (Biolegend), CD56 HCD56 (Biolegend), CD69 FN50 (Biolegend), CD71 CY1G4 (Biolegend), HLA-A,B,C W6/32 (Biolegend), IL-17 TC11-18H10.1 (Biolegend), IL-4 MP4-25D2 (Biolegend), IFN-y B27 (Biolegend), IL-22 2G12A41 (Biolegend), Granzyme A CB9 (Biolegend), Granzyme B QA18A28, TCR Vb18 BA62.6 (Beckman Coulter), TCR Vb7.1 ZOE (Beckman Coulter) and Beta Mark TCR Vbeta Repertoire Kit (Beckman Coulter). After staining, cells were washed twice with FACS buffer. Cells were then fixed and permeabilized using the Cytofix/Cytoperm kit (BD Biosciences) according to the manufacturer’s instructions. Intracellular HIV p24 Gag protein was stained using anti–HIV p24 antibody (clone KC57-RD1 or KC57-FITC, Beckman Coulter; clone 28B7, MediMabs) diluted in Perm/Wash buffer. The combination of these two antibodies targeting different HIV-Gag epitopes has been shown to improve the specificity and sensitivity of detection^51^. Cells were incubated for 30 minutes at room temperature in the dark, washed twice with Perm/Wash buffer, and resuspended in FACS buffer for acquisition. Samples were acquired on a BD Symphony and analyzed using FlowJo software (BD Biosciences).

### T-cell stimulation

To assess HIV reactivation, 100,000 cells per ARC and dARC were stimulated with T-cell activators anti-CD3/CD28 Dynabeads (Thermo Fisher) at 25 uL of beads per million cells or PMA (50 ng/mL) plus ionomycin (1 μg/mL) (PMA/I), in clone media. Stimulations were performed for 24 h at 37 °C with 5% CO₂. After stimulation, cells were stained with anti-HIV Gag antibodies conjugated to APC and RD-1, fixed, and analyzed by flow cytometry. HIV reactivation was quantified as the percentage of Gag+ cells, with dual staining used to identify latency reversal. Data were analyzed using FlowJo.

### Anti-CD3/CD28 stimulation to study latency reversal along with proliferation

Latently infected CD4⁺ T cells were cultured in clone media. To track proliferation, cells were labeled with CellTrace™ Violet (Thermo Fisher Scientific) according to the manufacturer’s protocol. Briefly, cells were incubated with 5 µM CellTrace Violet in PBS for 20 minutes at 37°C in the dark, followed by quenching in R10 medium and a 5-minute incubation at room temperature. Cells were plated at 1 × 10⁶ per well in U-bottom 96-well plates in clone media and either left unstimulated or stimulated with 1 µg/mL of soluble Ultra-LEAF™ purified anti-human CD3 (clone OKT3, BioLegend, Cat. No. 317325) and anti-human CD28 (clone 28.2, BioLegend, Cat. No. 302933) antibodies. Irradiated allogeneic PBMCs (from fresh buffy coat, irradiated at 30 Gy) were added as feeder cells at a 1:1 ratio. Cells were cultured for 4 and 7 days at 37°C with 5% CO₂, with media replenished as needed. At the end of culture, cells were harvested, fixed, and permeabilized using the Cytofix/Cytoperm kit (BD Biosciences) according to the manufacturer’s instructions. Intracellular HIV Gag was detected using APC-conjugated anti–HIV p24 antibody (clone 28B7 MediMabs) diluted in Perm/Wash buffer. Cells were incubated for 30 minutes at room temperature in the dark, washed twice with Perm/Wash, and resuspended in FACS buffer. Samples were acquired using a BD Symphony flow cytometer and analyzed with FlowJo software (BD Biosciences). Proliferation was assessed by dilution of CellTrace Violet, and latency reversal was quantified as the frequency of p24⁺ cells.

### HIV RNA quantitative PCR

Copies of HIV RNA was measured in 30 ml of cell culture supernatant using the previously described integrase single-copy assay protocol with minor modifications^64^. Briefly, QIAcube HT automated system (Qiagen, USA) and QIAamp 96 Viral RNA Kit-10 (Qiagen, USA) were used for RNA extraction and elution in 80 ml of water. A validated RNA standard was used for quantification of RNA samples. 12.5 ml of 2X buffer and 1 ml of N2 polymerase from AgPath-ID™ One-Step RT-PCR Reagents kit (ThermoFisher, USA) was mixed with 8.5 ml of extracted RNA or serially diluted RNA standards, 1 ml of 10 mM forward and reverse primers, and 1 ml of 6.25 mM probe, in a total volume of 25 ml per well. Samples and standards were run on each 96-well plate using a QuantStudio Pro 7 Real-Time PCR System (ThermoFisher Scientific). The cycling parameters were 50°C for 10 min, 95°C for 10 min, followed by 40 cycles of 95°C for 15 Sec and 60°C for 1 min. Cycle threshold values of samples and standards were analyzed by Design & Analysis Software v. 2.8.0 (ThermoFisher Scientific) to determine HIV RNA concentration. Primers and probes used in this assay are given in Supplementary Table 3.:

### Competitive ARC / dARC / newly infected cell killing assay

This assay was designed to parse viral from host factors in the susceptibility of resistance of an ARC to killing by HIV-specific CTL. Quality control of ARCs and dARCs: The day prior to an assay these were assessed by flow cytometry staining for the TCR Vb segment of a given ARC or dARC to confirm >95% purity. This was corroborated by IPDA. ARCs were stained and analyzed by flow cytometry for intracellular Gag ensuring sufficient spontaneous production to yield a readout. Generation of ARC Chr 16 viral stock: ARC Chr16 was washed 3x and resuspended in R10-100 at 1-2x10^6^ cells/mL in 4 mL for 24 hours. Supernatants were harvested, and virus was concentrated by PEG-it, following manufacturer’s instructions. Virus was frozen at -80°C in 5 x 50 mL aliquots and two aliquots were used for the infection. Preparation of target cells: CD4^+^ T-cells were enriched from donor OM5334 by negative selection using EasySep™ Human Memory CD4^+^ T Cell Enrichment Kit (STEMCELL Technologies) and plated at 5x10^6^ cells/ml in R10-100 (no ARVs) along with ImmunoCult™ Human CD3/CD28 T Cell Activator at 25 µL/mL (STEMCELL Technologies). After 48 hours, these cells were infected by adding the ARC Chr16 viral stock and incubating at 37°C for 2 hours. Cells were then washed with R10-100, plated at 5x10^6^ cells/ml in fresh R10-100 and incubated for 3 days, at which point they were harvested for the competitive killing assay. These are termed newly *in vitro* infected cells. Labeling of target cells: On the day of co-culture dARCs, ARCs, and newly *in vitro* infected cells were each washed and then labeled separately with different concentrations of CellTrace™ Yellow (CTY; Thermo Fisher Scientific) according to the manufacturer’s instructions. Newly *in vitro* infected cells were labeled with 5 uM CTY, ARC Chr16 with 10x less 5 uM CTY, and the dARC treated identically but without CTY. Small aliquots of each were analyzed by flow – separately and mixed – to ensure that staining intensities could be distinguished. Quality control of HIV-specific CTL clone: CTL clone was taken 3 weeks after its most recent restimulation cycle. Specificity and functionality was confirmed by an overnight ‘specificity check (see above) using the cognate HIV-Pol peptide. Co-culture: Labeled targets and CTL were co-cultured with each other at a range of effector:target ratios ranging from 1:2 to 1:32 a no CTL control in triplicate – approximately 2x10^5^ total cells/well in a 96-well plate in clone media, along with 1/100 anti-CD107a APC-Cy7 (Biolegend) to allow assessment of degranulation. Co-cultures proceeded overnight at 37°C. Cells were then surface stained with the following antibodies: MHC-I PeDazzle, CD4 alexa700, CD3 BV785, CD8 BV605 (all from Biolegend) and live/dead aqua (Thermo Fisher). Cells were fixed and permeabilized with the Cytofix/Cytoperm kit (BD Biosciences) and then labeled intracellularly for: HIV-Gag (KC57, Beckman Coulter), HIV-Gag (28B7, MediMabs), BCL-2 BV421 (Biolegend), and CD8 BV605 (re-labeled for internalized CD8). Cells were then fixed with PFA and analyzed on an Attune NxT flow cytometer.

### Enrichment / depletion of ARCs based on CD4 expression

ARC cells were stained with 20 uL of anti-human CD4 microbeads (Milteny Biotec) per 1x10^7^ cells in 80 uL of MACS buffer at 4°C for 15 minutes. Cells were then washed in 1 mL of MACS buffer and centrifuged at 400 x g for 5 minutes. Stained cells were resuspended in 500 uL of MACS buffer for up to 1x10^7^ cells. 500 uL of cells were loaded unto an MS MACS column on a miniMACS magnetic separator on a MACS multi stand magnet (Milteny biotec). Cell were washed on the magnet 3X with 500 uL of MACS buffer and flow through was collected as Gag+ enriched cells (CD4+ depleted). The cells bound to the column were then removed from the magnet and flushed with 1 mL of MACS using a plunger into a collection tube as Gag+-depleted cells (CD4+ enriched). Prepared cell fractions were then analyzed at day 0 by flow cytometry for Gag expression and also analyzed at day 5 and day 10 after being plated at 1 × 10⁶ per well in U-bottom 96-well plates in clone media and stimulated with 1 µg/mL of soluble Ultra-LEAF™ purified anti-human CD3 (clone OKT3, BioLegend) and anti-human CD28 (clone 28.2, BioLegend) antibodies.

### 5’ CITE-seq compatible with a 10X genomics instrument

Cells were stained with DNA oligo-barcoded antibodies and loaded onto the 10x Chromium single-cell workflow (10X Genomics) as previously described^65^ with a few modifications. Approximately 0.05-0.3 x 10^6^ cells per sample were resuspended in R10 media and incubated with TotalSeq-C Human Universal Cocktail V1.0 (Biolegend) for 30 minutes at 4°C. In some cases, TotalSeq™-C hashing antibodies TotalSeq™-C0251 anti-human Hashtag 1 clones LNH-94, 2M2 (Biolegend), TotalSeq™-C0251 anti-human Hashtag 3 clones LNH-94, 2M2 (Biolegend), TotalSeq™-C0251 anti-human Hashtag 4 clones LNH-94, 2M2 (Biolegend) were spiked into individual samples, which allowed for multiplexing of ARCs and uninfected clone combinations into the same 10X lane. After staining, cells were washed 3 times in R10 media, followed by centrifugation (400 × g for 5 minutes at 4°C) and supernatant aspiration. After the final wash, cells were resuspended in PBS and filtered through 70-µm cell strainers. Stained cells from each sample were pooled and loaded into the 10x Chromium Single Cell Immune Profiling workflow (CG000330, revision G). Libraries were generated according to the 10x Chromium Single Cell Immune Profiling workflow instructions, pooled and sequenced on an Illumina NovaSeq 6000 or NovaSeqX.

### Single-cell data alignment

FASTQ files from the 10x gene expression, surface protein and TCR α/β libraries were processed using Cell Ranger v7.0.1 or v9.0.1 (10x Genomics), using either the count or the multi pipeline (when TCR libraries were available) with default parameters.

RNA transcripts from ECCITE-seq were mapped to a modified version of Cell Ranger’s GRCh38-2020-A reference package (GRCh38 / GENCODE v32 annotations), with the addition of the HIV HXB2 reference sequence. TCR α/β libraries were mapped to Cell Ranger’s cellranger-vdj-GRCh38-alts-ensembl-7.1.0 reference package.

### Initial cell/gene level filtering

For cell filtering, cell barcodes with <200, >8,000 genes expressed or > 10% mitochondrial genes expressed were removed. For gene-level filtering, genes were filtered such that only those expressed at a level of <200 UMI or in < 3 cells were removed.

These cell- and gene-level filtered gene/barcode matrices were used for downstream analysis.

### Seurat normalization

The count matrices of all modalities were loaded into a Seurat v5 object (Seurat version 5.1.0). For gene expression (GEX) data, we normalized the count data using LogNormalize and scaled the resulting normalized values using ScaleData. For all other modalities (ADT, HTO, TCR α/β), we normalized the count data using centered-log ratio (margin = 2) and scaled the resulting normalized values using ScaleData.

### Hashing antibody demultiplexing

To separate individual samples from pooled cell libraries, samples were demultiplexed by their HTO using Seurat HTODemux with the positive quantile parameter set to 0.999.

### Multimodal WNN analysis

We applied weighted-nearest neighbor (WNN) analysis on our 5’CITE-seq data, enabling the integrative analysis of GEX and ADT modalities in the same cells as previously described^20^. Briefly, k-nearest neighbors graphs (k = 20) were first constructed, and the neighbors of each cell in each modality were identified using FindMultiModalNeighbors in Seurat V5. For graph construction, the first 15 principal components (PCs) of the log-normalized values from the GEX modality and the first 12 PCs of the centered-log ratio values from the ADT modality were used. The dimensionally reduced molecular profiles of the k-nearest neighbors of each cell were then averaged and compared with the actual values to obtain the residuals in an RNA-RNA, protein-protein and RNA-protein manner. Residuals were converted into affinity-based similarities using UMAP. Ratios of the within-modality and cross-modality affinities were softmax transformed to obtain the cell-specific modality weights. The weighted similarity then formed the basis of the construction of the WNN graph visualized on a WNN integrated UMAP. Clusters were called using the FindClusters function at a resolution of 0.5.

### Differential gene expression and pathway enrichment analysis

Differential gene expression analysis across WNN clusters was performed using FindAllMarkers on cells defined as ARCs or uninfected clones based on hashing information and TCR sequences. The logfc.threshold (minimum log_2_ fold change) was set to 0.1 and min.pct (minimum proportion of cells expressing the given gene) was set to 0.01. Only genes with an adjusted P-value of < 0.05 were further considered. Row normalized Z-scores produced using scale.data were used for visualization on the heatmap focusing on the top 15 differentially expressed genes.

Genes passing the above filters were used for pathway enrichment analysis. ClusterProfiler was used to profile clone populations on Reactome pathways. Reactome terms were considered significant if the FDR-adjusted P-value was < 0.05.

### Module scoring on ARCs and uninfected clones

The AddModuleScore function of Seurat V5 was used for scoring gene expression modules. For Reactome or Gene Ontology gene-sets, the msigdbr package was used to pull selected gene-sets from the Molecular Signatures Database (MSigDB).

As above, the AddModuleScore function of Seurat V5 was used for scoring gene expression module of the “CD8 cytotoxic” functional state gene-set from Szabo *et* al., 2019^43^

### Generation and maintenance of HIV-specific CTL clones

CTL clones were generated as previously described^66–68^. Peripheral blood mononuclear cells (PBMCs) from PWH were plated at 10^7^ cells/ mL in 1 mL of X-VIVO serum-free medium (Lonza) supplemented with HIV peptide pools (NIH AIDS reagent program) at a final concentration of 10 µg/mL/peptide and incubated overnight. T-cells producing IFN-γ in response to peptide were enriched using the Miltenyi Biotec IFN-γ Secretion Assay – Cell Enrichment and Detection Kit (PE) (Cat. No. 130-054-201), following manufacturer’s instructions. Cell separation was carried out using the OctoMACS™ magnet and MACS® MS Columns (Miltenyi Biotec). Enriched cells were subjected to four 5x serial dilutions in 10mL R10-50-15 each plated on a full 96-well plate. Feeder media was added containing 1x10^6^ irradiated (5,000 rad) PBMCs / mL from fresh buffy coats (HIV-negative donors), along 100 ng/mL each of Ultra-LEAF™ purified anti-human CD3 antibody (clone OKT3, BioLegend, Cat. No. 317325) and Ultra-LEAF™ purified anti-human CD28 antibody (clone 28.2, BioLegend, Cat. No. 302933). After 3 weeks, wells from the lowest dilution plate showing growth were screened for responsiveness to the target peptide using the ‘CTL specificity check’ (see below).

### CTL specificity check

CTLs were assessed for peptide specificity four weeks after isolation. Cells were resuspended in R10-50 medium supplemented with 10 ng/mL IL-15 and plated into 96-well plates. To each well, 2 µL of anti-CD107a (LAMP-1) PE-conjugated antibody (BioLegend) was added. The cognate peptide was added to half of the wells at a final concentration of 10 µg/mL. All wells were brought to a final volume of 200 µL with media. Cells were incubated at 37°C for 4 hours to allow for degranulation marker expression. Following stimulation, cells were stained with a surface antibody master mix composed of 0.5 µL anti-CD3 (BioLegend), 0.5 µL anti-CD4 (BD Biosciences), 0.5 µL anti-CD8 (BioLegend), 0.3 µL of viability dye (Thermo Fisher Scientific), and 48.2 µL of MACS buffer per well. Plates were covered and incubated at room temperature for 20 minutes, then washed twice with MACS buffer. Cells were fixed in 4% paraformaldehyde (VWR) and analyzed by flow cytometry to determine antigen-specific CD8⁺ T cell degranulation.

### SLOW CURE Assay

Authentic HIV reservoir clones were co-cultured with autologous HIV-specific cytotoxic T lymphocytes (CTLs) at a 1:1 ratio in R10 medium (RPMI 1640 supplemented with 10% FBS, 1% penicillin/streptomycin, 1% HEPES, and 1% L-glutamine). Prior to co-culture, reservoir clones were labeled with CellTrace™ Violet (CTV; Thermo Fisher Scientific) according to the manufacturer’s protocol. Briefly, CTLs were incubated with 5 µM CTV in PBS for 20 minutes at 37°C in the dark, followed by quenching with five volumes of R10 medium and a 5-minute incubation at room temperature. Irradiated allogeneic feeder cells were prepared from freshly isolated PBMCs obtained from healthy donors and irradiated with 30 Gy using a gamma irradiator to prevent proliferation. These feeder cells were added to the co-culture at a 1:1 ratio with the total number reservoir clones only or of CTLs and reservoir clones combined in duplicate conditions for comparison. Technical duplicates for each condition were set up at the start of culture Co-cultures were maintained in U-bottom 96-well plates at 37°C with 5% CO₂ for 1, 5 and 7 days for readout. Supernatant from culture were also harvested for viral load measurements. Media was replenished as needed to maintain cell viability. In some cases reservoir clones alone or in combination with CTLs were treated with 10 µM of Deferoxamine (DFO) in CTL sensitization experiments.

At the end of the incubation period, cells were harvested and spiked with CountBright™ Absolute Counting Beads (Thermo Fisher) and stained for surface markers including TCRvb specificity when available to more accurately track clonal dynamics in culture and intracellular HIV p24 Gag protein using APC-conjugated anti-p24 antibody (clone 28B7; NIH HIV Reagent Program, Cat. No. ARP-1475) in combination with a FITC-conjugated anti-p24 antibody (clone KC57; Beckman Coulter, Cat No. 6604665. Staining was performed following fixation and permeabilization with the eBioscience™ Foxp3 / Transcription Factor Staining Buffer Set (Thermo Fisher) according to manufacturer’s instructions. Proliferation of CTLs was assessed by dilution of CTV fluorescence intensity using flow cytometry (BD Symphony A5), and the total counts of reservoir clones and p24⁺ cells as well as p24⁺ frequency were quantified to evaluate clone propagation and survival and viral expression. Data analysis was performed using FlowJo software (BD Biosciences).

### In vitro infection of memory CD4^+^ T-cells

Memory CD4^+^ T cells were magnetically isolated from thawed PBMCs using the EasySep™ Human memory CD4^+^ T Cell Isolation Kit (Stemcell) according to the manufacturer’s instructions. Purified memory CD4^+^ T cells were then activated using anti-human CD3/CD28-coated magnetic beads (one bead for two cells, Gibco Cat. #11131D) in R10 media. After 3 days, cells were maintained at a concentration of 10^6^ cells /mL in media containing 50 IU of human recombinant IL-2. At day 7, the cultured memory cells were infected with HIV JRCSF. HIV JRCSF plasmid was obtained from BEI-resources and virus was produced as previously described^56^. One quarter of the cells were infected by the addition of 200 units of virus per 1x10^7^ cells in 1mL and centrifugation at 1500xg for 2h at 37°C. Following this ’spinoculation’, the infected cell cultures were mixed in flasks containing the remaining three quarters of the cells at 2x10^6^ cells/mL in media containing 50 IU of human recombinant IL-2. At day 10, the cells were crowded in 96-well round bottom plates at a density of 2x10^5^ cells/200uL/well to enhance cell-to-cell transmission of HIV. On day 13, cells were cryopreserved in FBS with 10% DMSO and stored in liquid nitrogen for future experiments.

### Killing assays of *In vitro*-infected memory CD4^+^ T-cells

Infected memory CD4^+^ T-cells were thawed and rested for 2 days at 2 x 10^6^ cells per mL in media containing 50 IU of human IL-2. Prior to co-culture with CTL, cells expressing high levels of CD4 were depleted by magnetic selection using the EasySep™ Human CD4 Positive Selection Kit II (Stemcell); this step enriches for CD4^-^ infected cells (since HIV-infected cells downregulate CD4). Cells infected with HIV JRCSF were labeled with cell trace far red (CTFR) (dilution 1:20,000). TW10-Gag specific CTL were left unlabeled and were added to each well at a 1:2 effector:target ratio along with anti-CD107a APC-Cy7 (dilution 1:200). Killing assays were performed in R10 media supplemented with 50 IU of human recombinant IL-2 and 0.51 ng/mL of human recombinant IL-15 for 16h at 37°C.

### Intracellular cytokine staining of ARCs and uninfected clones

To confirm functional T-helper subtypes amongst ARCs and uninfected clones, cells were reactivated for 4 hours with the eBioscience™ Cell Stimulation Cocktail (plus protein transport inhibitors) (ThermoFisher) following the manufacturer’s instructions. Cells were stained with surface antibodies targeting CD4 (OKT4, Biolegend), CD8 (SK1, Biolegend), CD71 (CY1G4, biolegend), CD69 (FN50, Biolegend), along with a fixable live/dead stain (ThermoFisher) in PBS for 30 minutes at 37°C. Cells were washed and fixed with 1X eBioscience™ FoxP3/transcription factor fixation/permeabilization solution (ThermoFisher) for 45 minutes at 4°C. This fixation solution was washed away and cells were stained intracellularly with anti-HIV-Gag-p24 (KC57, Beckman Coulter), anti-IL-17 (TC11-18H10.1, Biolegend), anti-IL-4 (MP4-25D2, Biolegend), anti-IFN-γ (B27, Biolegend) and anti-IL-22 (2G12A41, Biolegend) antibodies in 1X FoxP3/transcription factor permeabilization solution (ThermoFisher) for 30 minutes at 37°C. Cells were washed one more time before analysis on an Attune NxT flow cytometry instrument.

## RESOURCE AVAILABILITY

### Data availability

All data necessary to evaluate and understand the conclusions of this manuscript are provided in this article or the supplementary information associated with it. Individual data points used for summary data display items are all included in the Source data provided with this paper. The proviral sequences reported in this article have been deposited in GenBank under accession numbers PX992673-PX992682. The 5’ CITE-seq data reported in this article has been deposited in the Gene Expression Omnibus (GEO) database and made publicly available under accession number GSE303197. Other data are available from the corresponding author upon reasonable request.

### Code availability

This paper does not report any original code. Code used for analysis of CITE-seq data will be made available upon request.

---

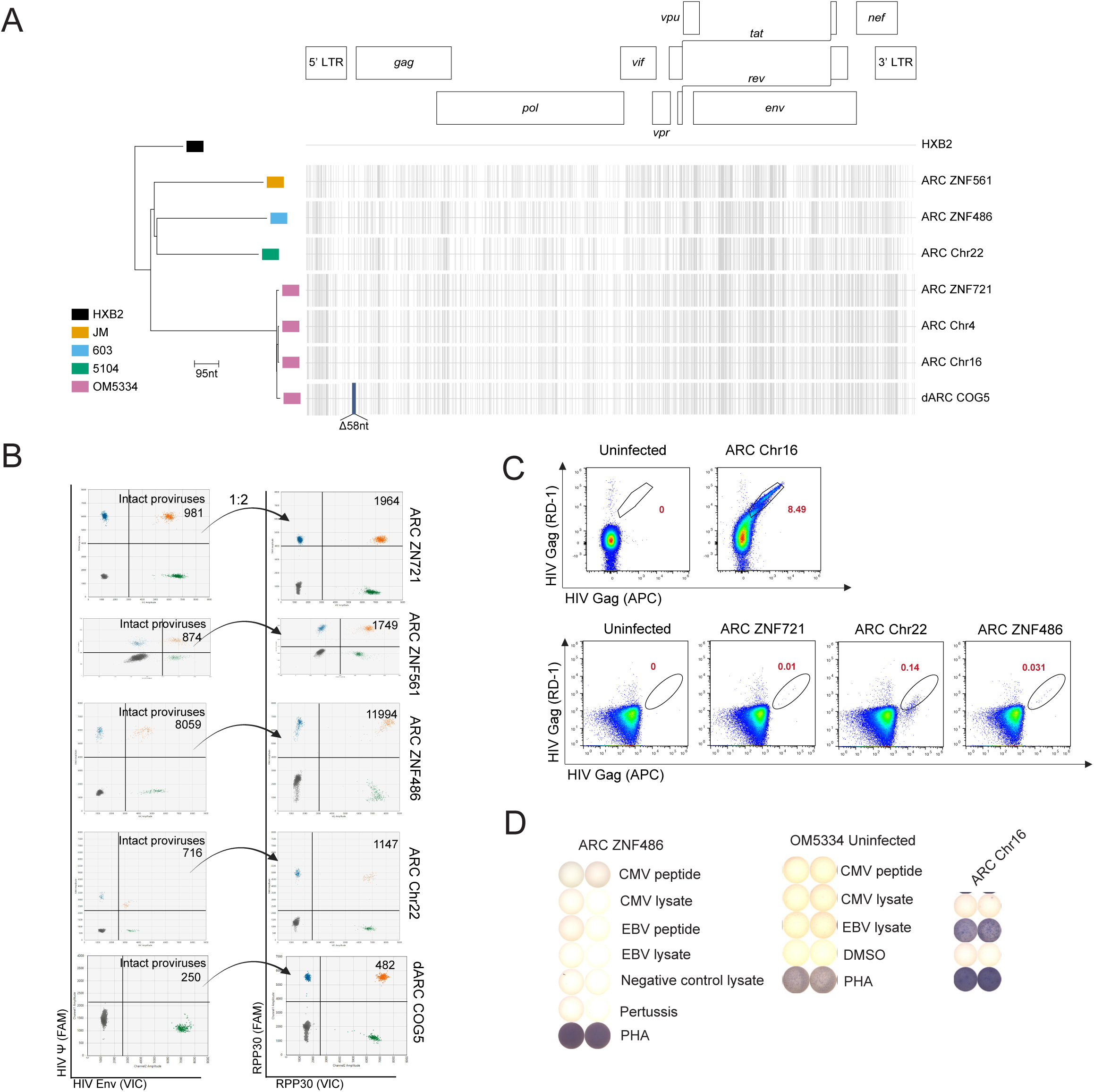

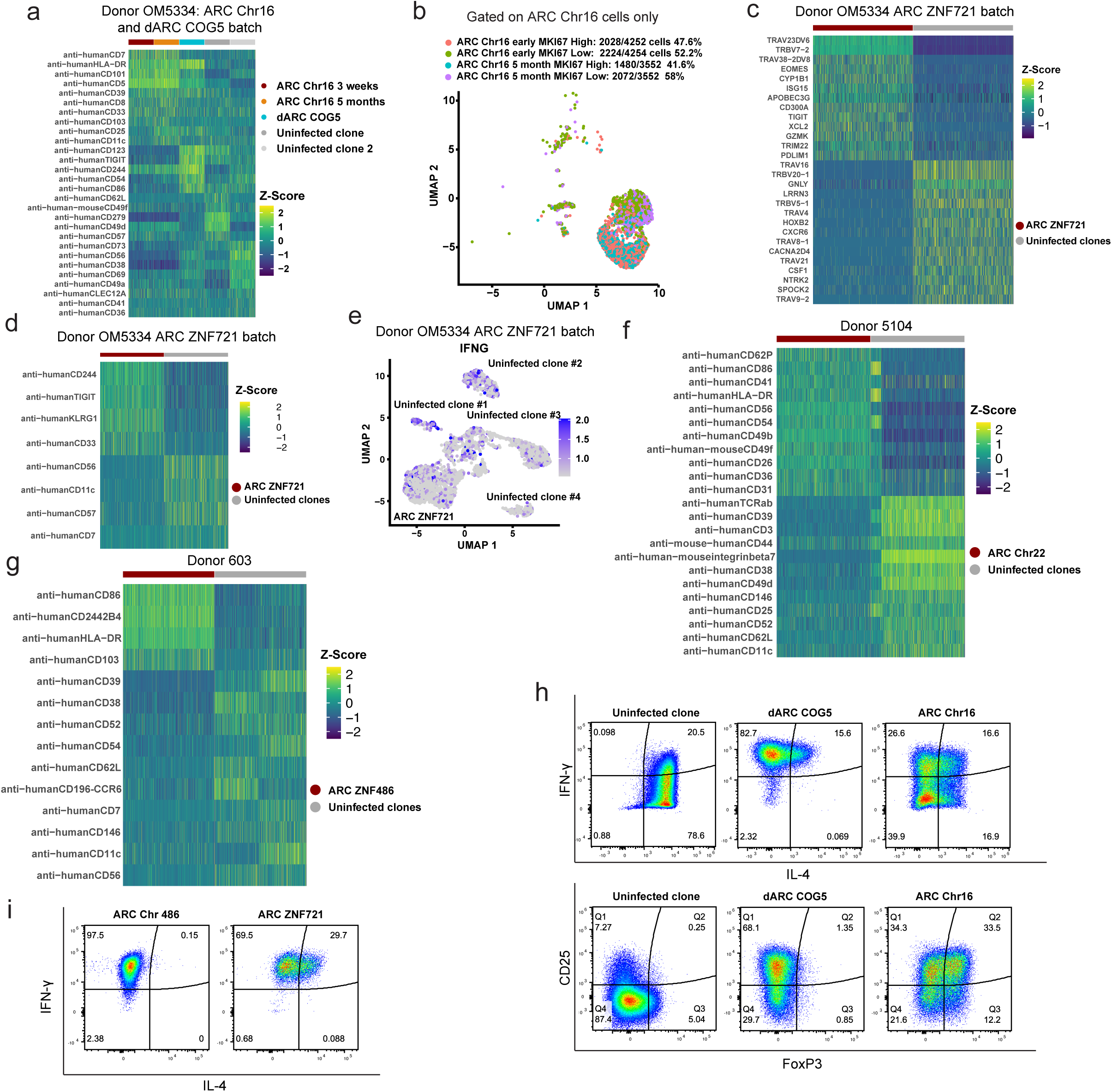

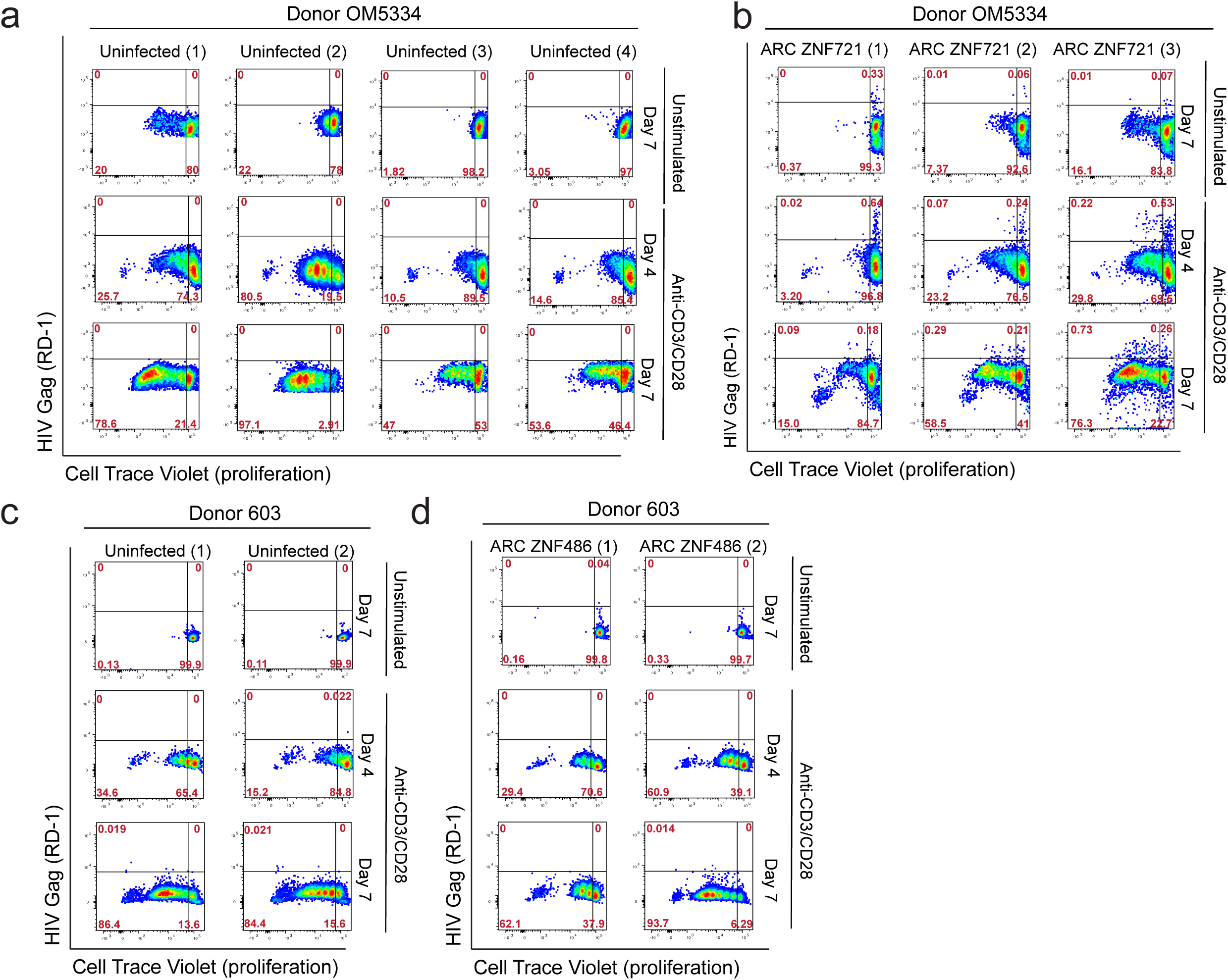

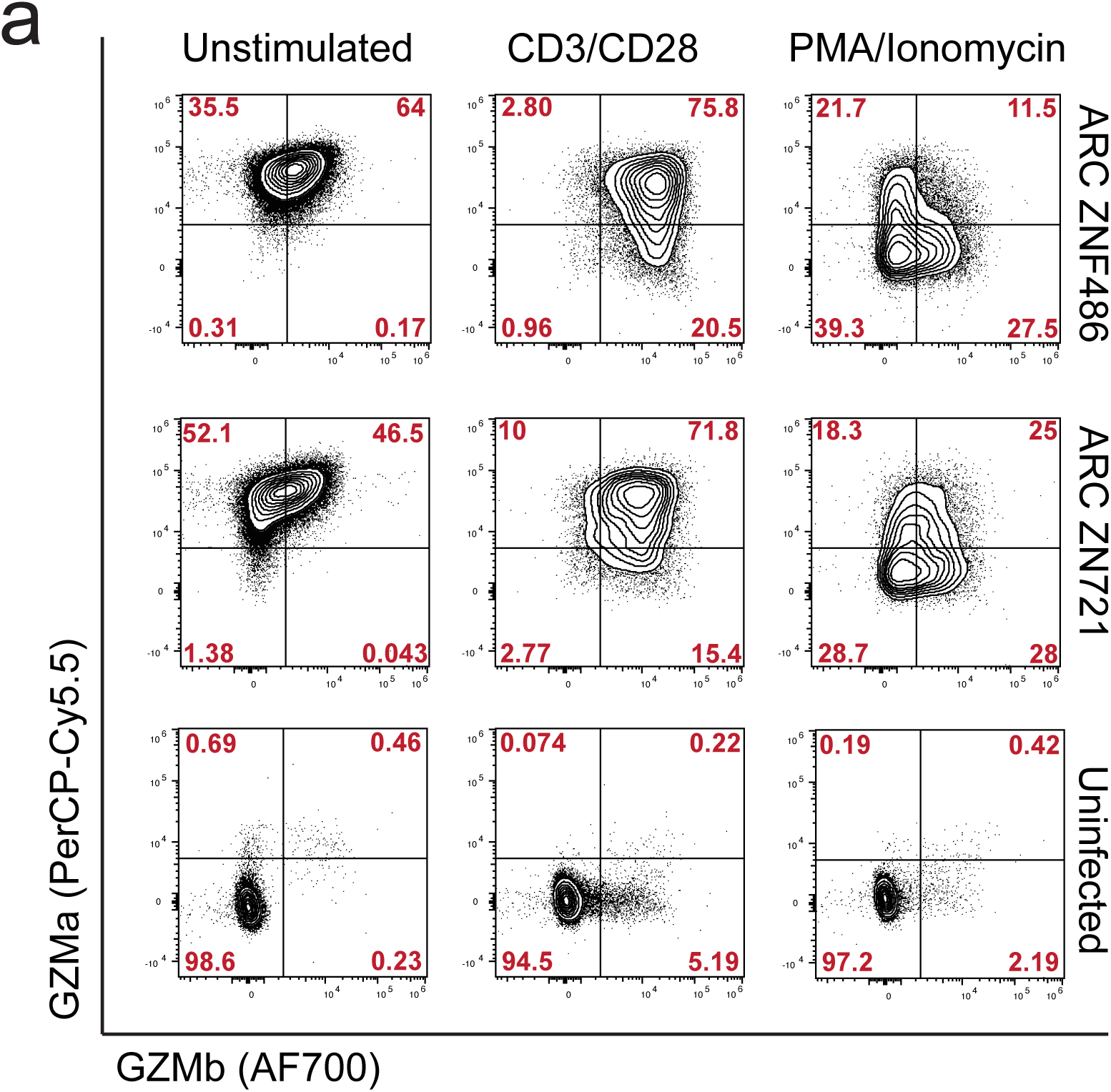

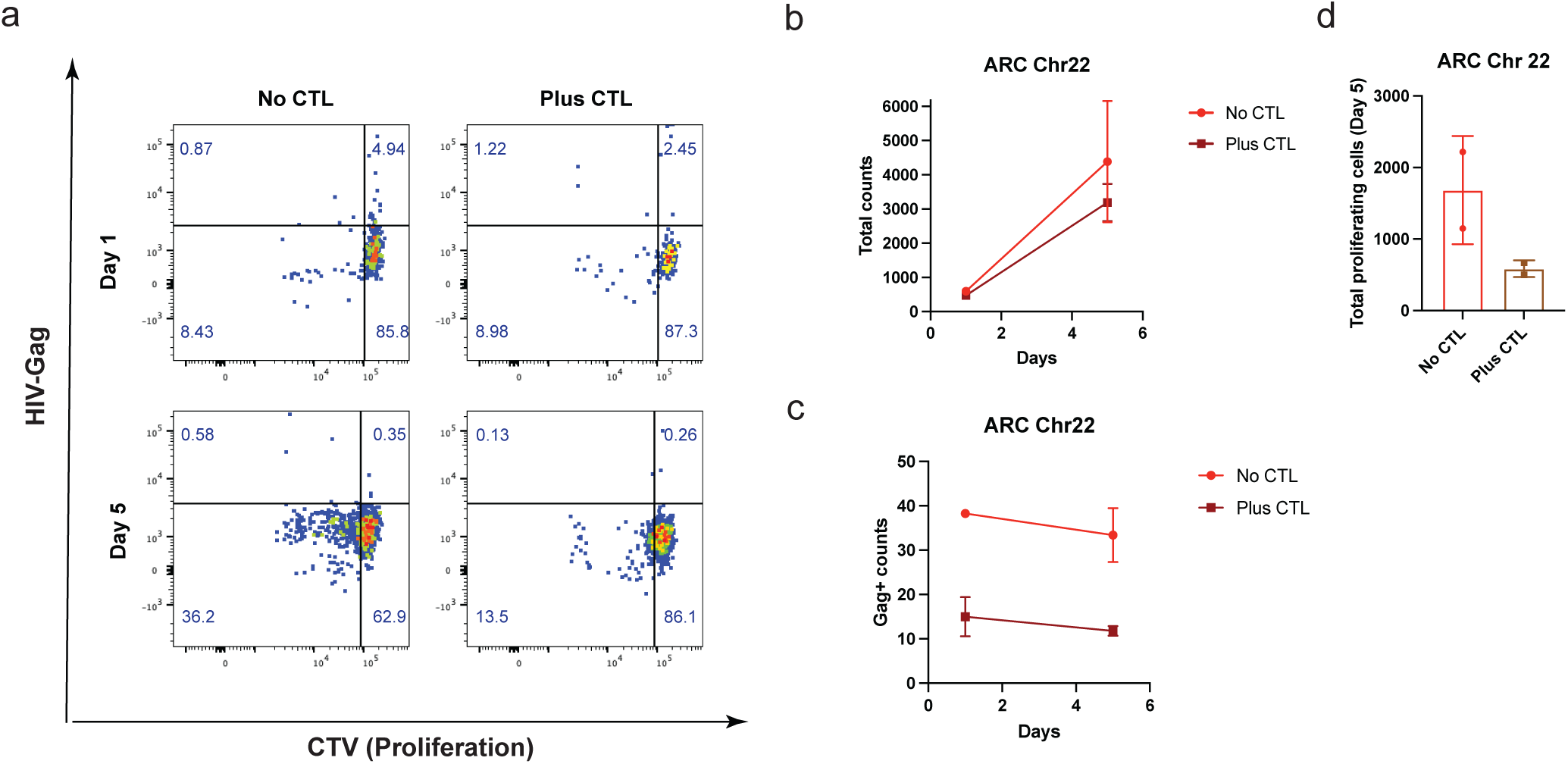

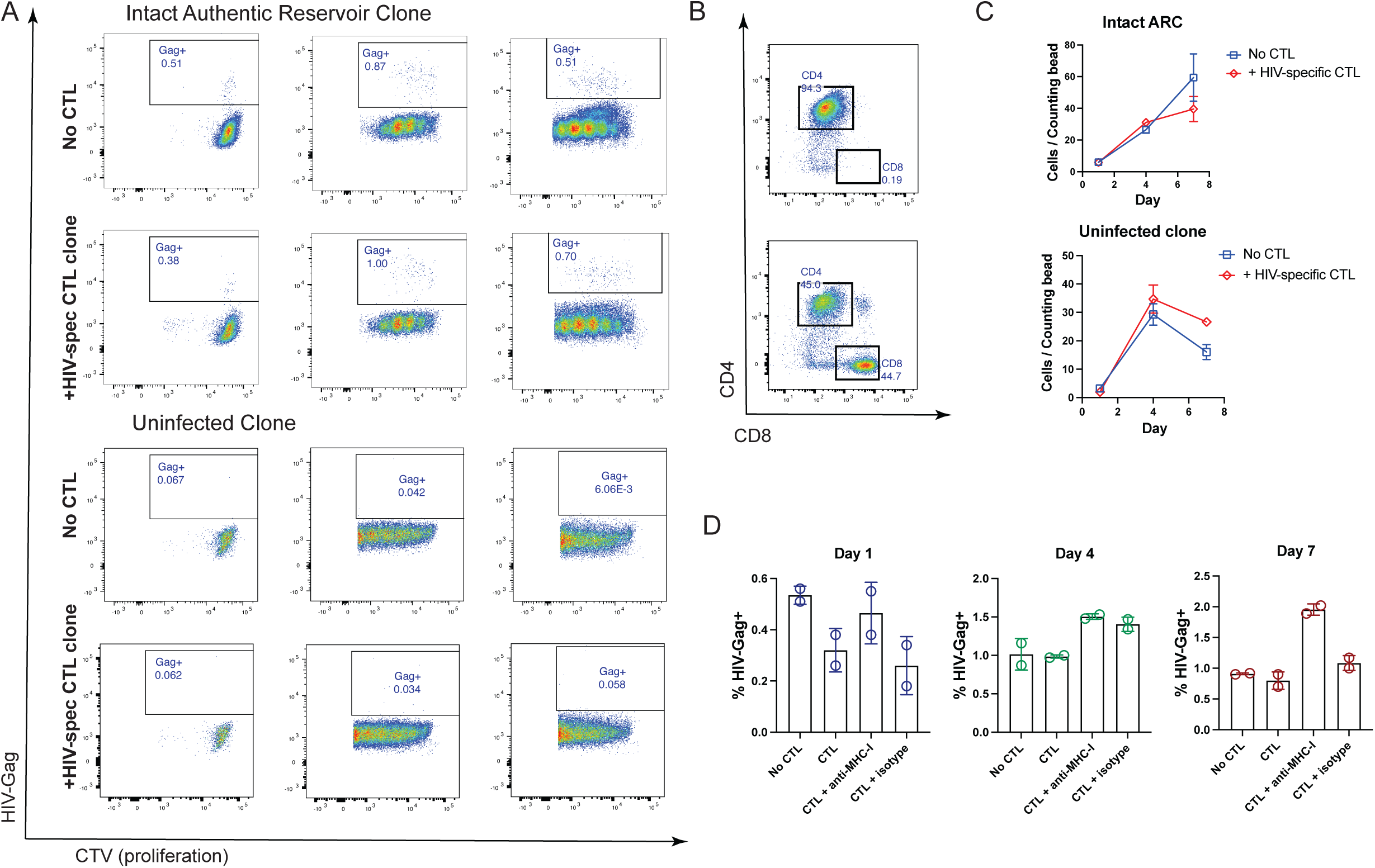

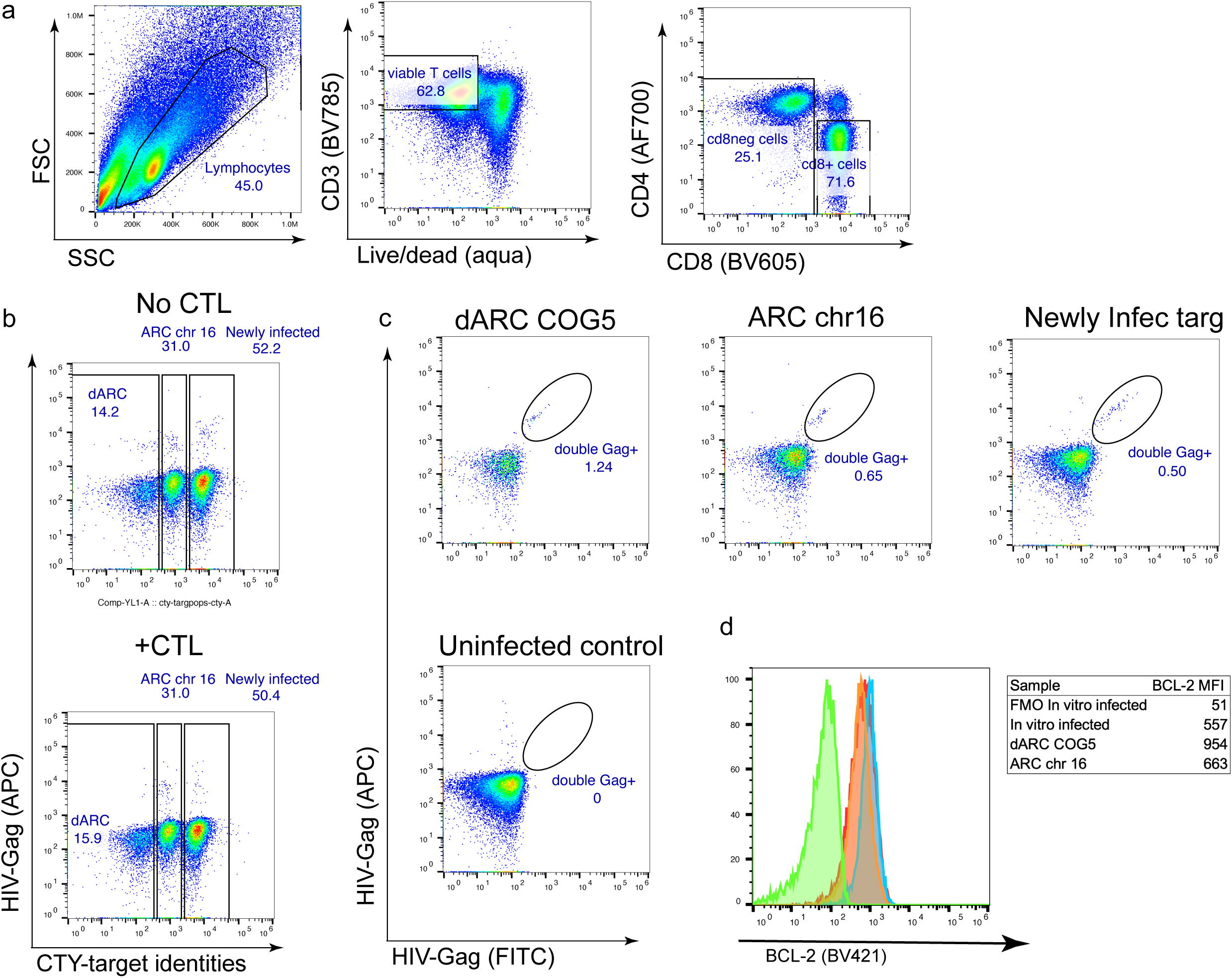

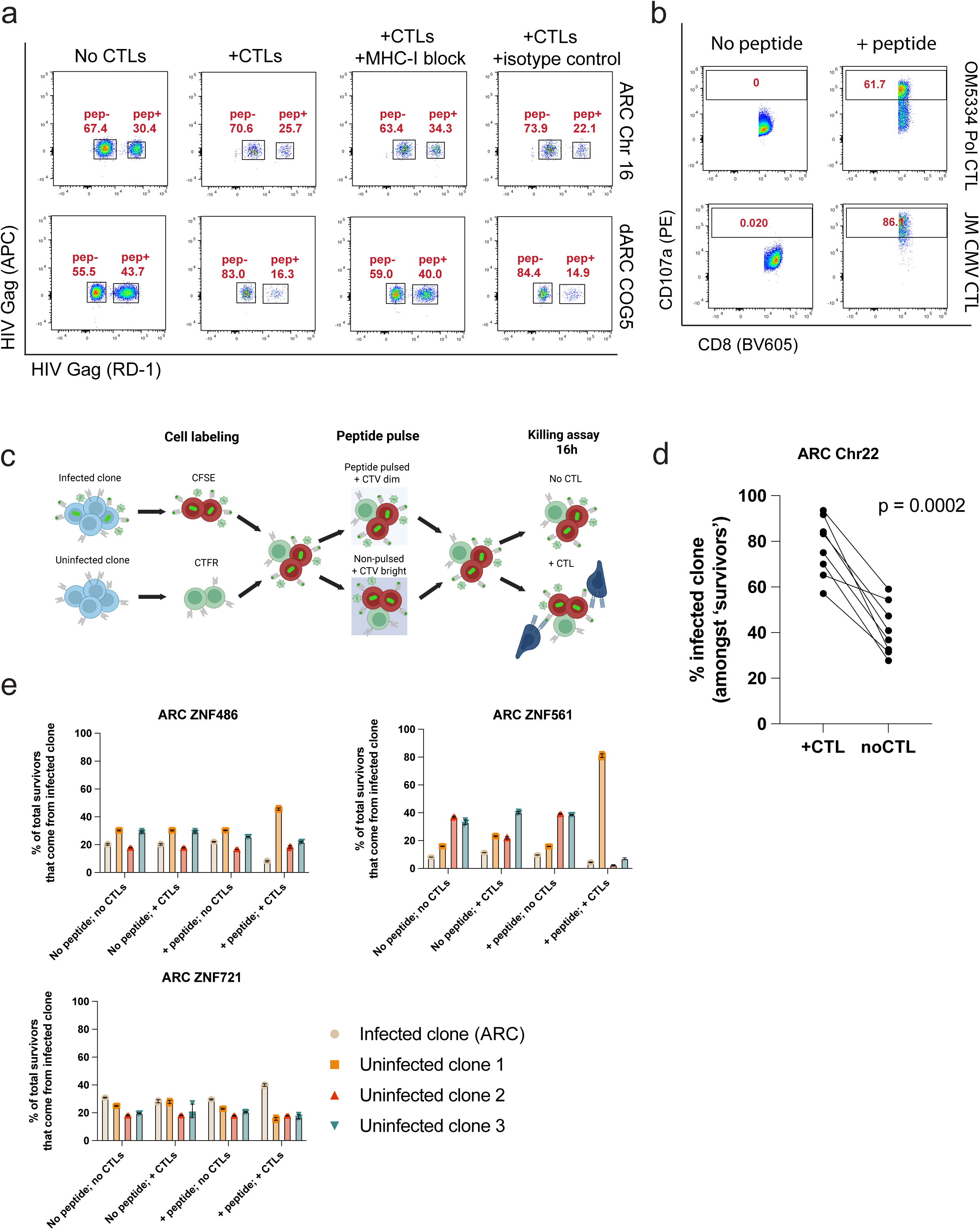

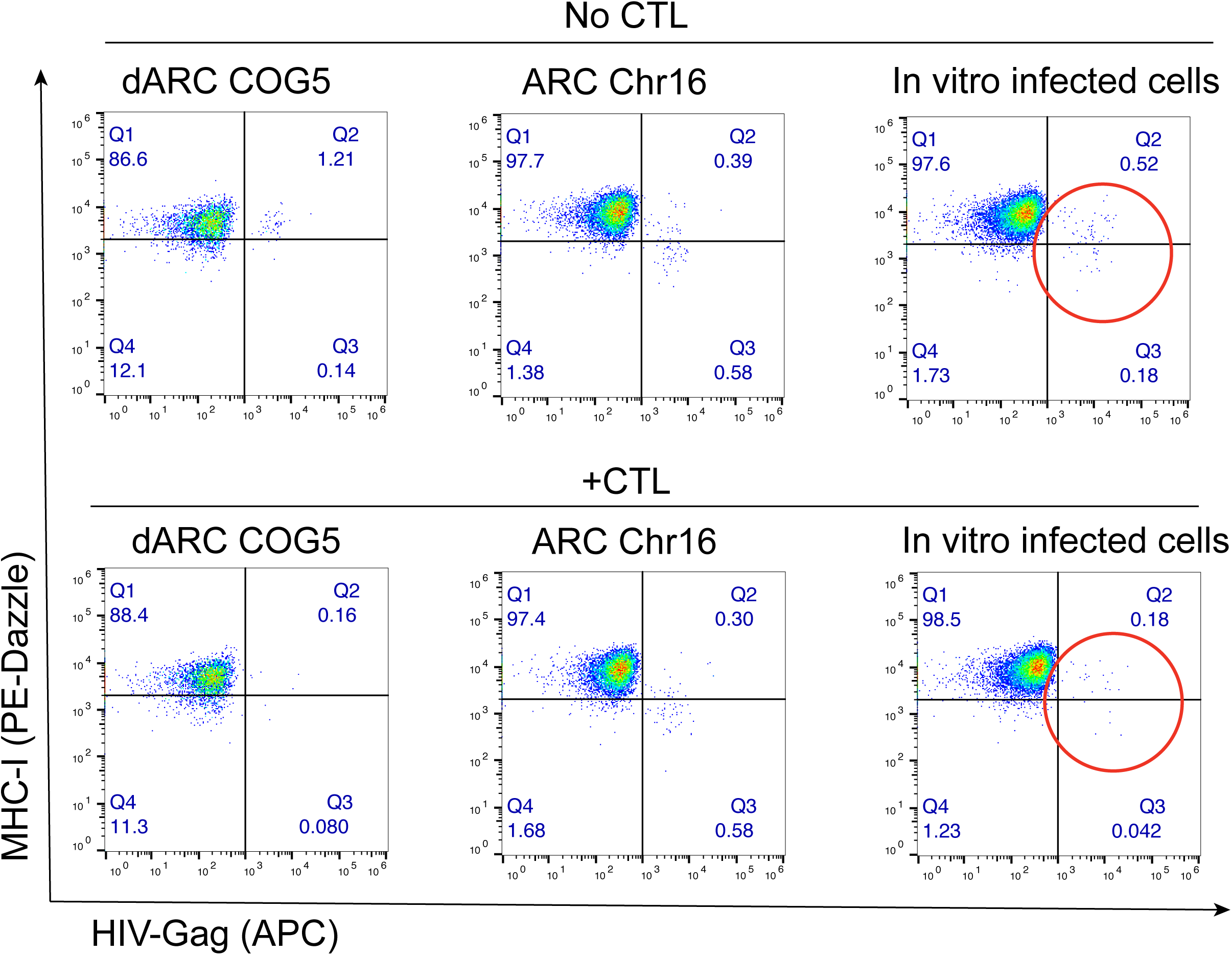

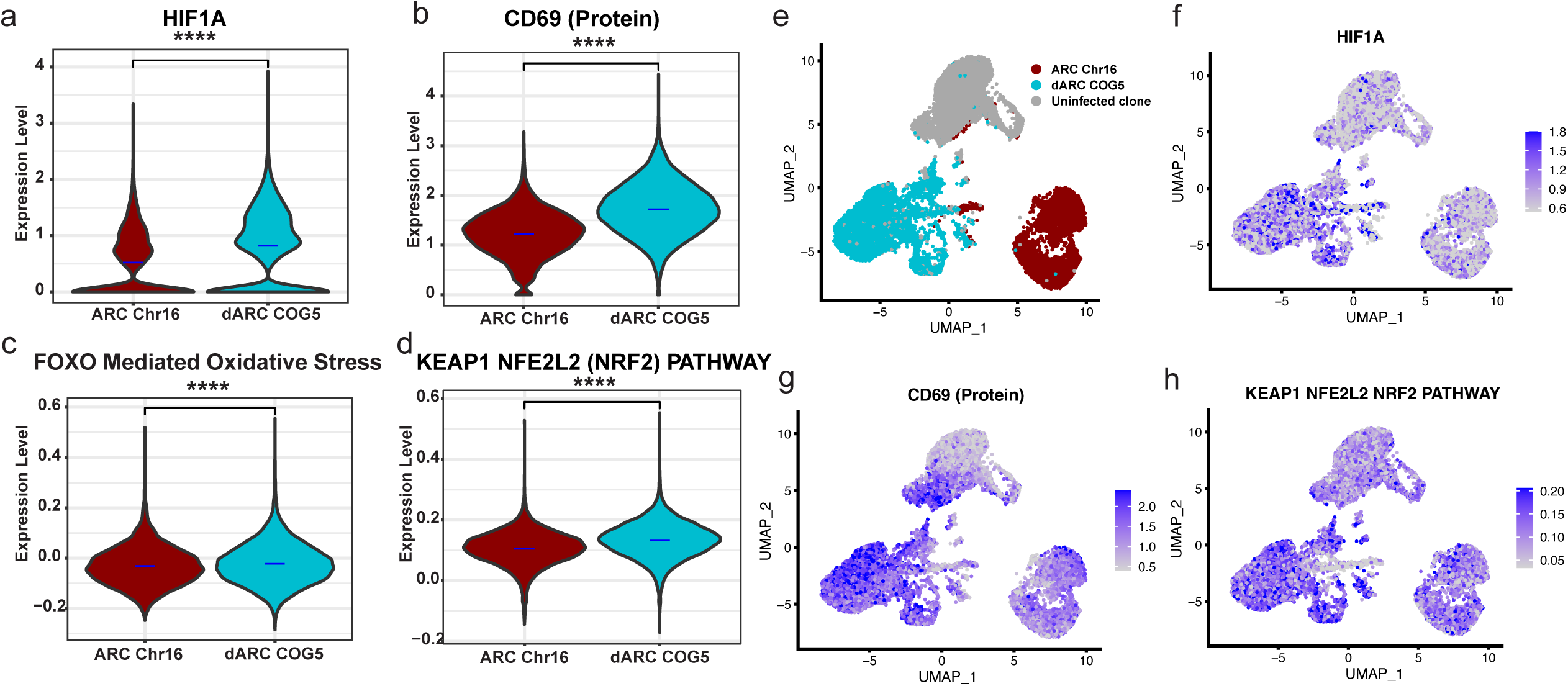

