## Supplementary tables for "Dynamic Antigen Expression and Intrinsic CTL Resistance in HIV Reservoir Clones"

**Supplementary Table 1.** Clinical and demographic data of donors.

| Donor ID | Age | Sex | Ethnicity | Years Since Diagnosis | ART Duration (yrs) | $CD4^+$ Nadir (cells/µl) | Current $CD4^+$ (cells/µl) | Plasma VL (copies/mL) | Suppression Duration (yrs) | ART regimen |
|---|---|---|---|---|---|---|---|---|---|---|
| 603 | 43 | M | White/Hispanic | 12 | 10 | 693 | 300 | <20 | 10 | EFV/TDF/FTC |
| 5014 | 35 | M | Black | 7 | 7 | 606 | 400 | <20 | 7 | BIC/TAF/FTC |
| OM5334 | 41 | M | White | 17 | 17 | 460 | 1016 | <20 | 10 | B/F/TAF |
| 908 | 57 | M | White | 7 | 5 | 314 | 484 | <40 | 5 | ABC/3TC/DRV/RTV |

**Supplementary Table 2.** ddPCR primer and probe sequences for ARC-specific quantitation.

| Assay | Primer type | Primer sequence |
|---|---|---|
| HIV-1 LTR (R-U5) | Forward | CTTAAGCCTCAATAAAGCTTGCC |
| | Probe | 5fam/AGTAGTGTG/zen/TGCCCGTCTG/3iabkfq |
| | Reverse | GGATCTCTAGTTACCAGAGTC |
| OM5334 ZNF721 integration site | Forward | GGAACTCAACGACAAATGTGTGTATACATATG |
| | Probe | 5hex/AGCCCTTCC/zen/ACACACACAAACAGAC/3iabkfq |
| | Reverse | GCCTTGTGTGTGGTAAATCCAC |
| OM5334 ZNF721 TCRβ CDR3 | Forward | TGACGATCCAGCGCACACAG |
| | Probe | 5cy5/CAGCTTAGA/tao/ATCAGGGTTCCATCAGC/3iabkrq |
| | Reverse | GGTCCTCTAGGATGGAGAGTCG |

**Supplementary Table 3.** ddPCR primer and probe sequences.

| **Primers and probes used for quantitation by digital droplet PCR** | | | |
|---|---|---|---|
| Assay | Oligo type | Oligo sequence | Reference |
| **RPP30 ddPCR** | Forward | GATTTGGACCTGCGAGCG | Stevenson EM et al. (35985993) |
| | Probe | VIC/CTGACCTGAAGGCTCT/MGBNFQ | |
| | Reverse | GCGGCTGTCTCCACAAGT | |
| | Forward | CCATTTGCTGCTCCTTGGG | |

| | | | |
|---|---|---|---|
| **RPP30-shear ddPCR** | Probe | FAM/AAGGAGCAAGGTTCTATTGTAG/MGBNFQ | |
| | Reverse | CATGCAAAGGAGGAAGCCG | |
| **HIV-1 Ψ ddPCR** | Forward | CAGGACTCGGCTTGCTGAAG | Bruner KM et al. (30700913) |
| | Probe | FAM/TTTTGGCGTACTCACCAGT/MGBNFQ | |
| | Reverse | GCACCCATCTCTCTCCTTCTAGC | |
| **HIV-1 env ddPCR** | Forward | AGTGGTGCAGAGAGAAAAAAGAGC | |
| | Probe | VIC/CCTTGGGTTCTTGGGA/MGBNFQ | |
| | Hypermutant competition probe | CCTTAGGTTCTTAGGAGC/MGBNFQ | |
| | Reverse | GTCTGGCCTGTACCGTCAGC | |
| **HIV-1 gag back-up ddPCR** | Forward | TCTCGACGCAGGACTCG | Stevenson EM et al. (35985993) |
| | Probe | FAM/CTCTCTCCTTCTAGCCTC/MGBNFQ | |
| | Reverse | TACTGACGCTCTCGCACC | |
| **HIV-1 env back-up ddPCR** | Forward | ACTATGGGCGCAGCGTC | Gramatica A et al. (40333189) |
| | Probe | VIC/CTGGCCTGTACCGTCAG/MGBNFQ | |
| | Reverse | CCCCAGACTGTGAGTTGCA | |
| | Forward | CTTAAGCCTCAATAAAGCTTGCC | Anderson EM, |

| | | | |
|---|---|---|---|
| **HIV-1 LTR (R-U5) ddPCR** | Probe | FAM/AGTAGTGTG/ZEN/TGCCCGTCTG/3IABkFQ | Maldarelli F (30253074) |
| | Reverse | GGATCTCTAGTTACCAGAGTC | |
| **603 ZNF486 integration site ddPCR** | Forward | TGCTGGGATTACAGGTGTGAG | Original to this study |
| | Probe | HEX/AATTAACCC/ZEN/TTCCACCAACAAAAGATCT/3IABkFQ | |
| | Reverse | GTAGCCTTGTGTGTGGTAGACC | |
| **603 ZNF486 TCRβ ddPCR** | Forward | TCCTCTCACTGTGACATCGGC | |
| | Probe | Cy5/AGCTCGACC/TAO/CCTGTCCCGGTCTACTAC/3IAbRQSp | |
| | Reverse | CTGGTCCCGTTCCCAAAGTG | |
| **OM5334 chr16 integration site ddPCR** | Forward | GAATCATCATTGAATGGAATCGAATGG | |
| | Probe | HEX/AGTGAATTA/ZEN/GCCCTTCCAATTACATTGG/3IABkFQ | |
| | Reverse | AATCCACAGATCTAGAATGTCTTGCC | |
| **OM5334 COG5 integration site ddPCR** | Forward | CCTATGCTCTTTACAAAGCAGGG | |
| | Probe | HEX/ATTAGCCCT/ZEN/TCCAACTATTTTAGGTA/3IABkFQ | |
| | Reverse | AATGTCTTGCCTTGTCTGGG | |
| **OM5334 ZNF721 integration site ddPCR** | Forward | GGAACTCAACGACAAATGTGTGTATACATATG | |
| | Probe | HEX/AGCCCTTCC/ZEN/ACACACACAAACAGAC/3IABkFQ | |
| | Reverse | GCCTTGTGTGTGGTAAATCCAC | |

| | | | |
|---|---|---|---|
| **OM5334 ZNF721 TCRβ ddPCR** | Forward | TGACGATCCAGCGCACACAG | |
| | Probe | Cy5/CAGCTTAGA/TAO/ATCAGGGTTCCATCAGC/3IAbRQSp | |
| | Reverse | GGTCCTCTAGGATGGAGAGTCG | |
| **Primers used for near-full-length HIV-1 sequencing** | | | |
| Assay | Oligo type | Oligo sequence | Reference |
| **FLIP-Seq outer PCR** | Forward | AAATCTCTAGCAGTGGCGCCCGAACAG | Lee GQ et al. (28628034) |
| | Reverse | TGAGGGATCTCTAGTTACCAGAGTC | |
| **FLIP-Seq nested PCR** | Forward | GCGCCCGAACAGGGACYTGAAARCGAAAG | |
| | Reverse | GCACTCAAGGCAAGCTTTATTGAGGCTTA | |
| **Primers used for HIV-1 integration site sequencing** | | | |
| Assay | Oligo type | Oligo sequence | Reference |
| **Integration site adaptor ligation** | Adaptor | GTAATACGACTCACTATAGGGCACGCGTGGTCGACGGCCCGGGCTGCT | Einkauf KB et al. (30688658) |
| | Adaptor | /Phos/AGCAGCCC/AmMO/ | |
| **Integration site 5' outer PCR** | Forward | GTAATACGACTCACTATAGGGC | |
| | Reverse | GCTTCAGCAAGCCGAGTCCTGCGTCGAG | |
| **Integration site 5' nested PCR** | Forward | ACTATAGGGCACGCGTGGT | |
| | Reverse | GCTCCTCTGGTTTCCCTTTCGCTTTCAA | |

| | | |
|---|---|---|
| **Integration site 3' outer PCR** | Forward | CTTAAGCCTCAATAAAGCTTGCCTTGAG |
| | Reverse | GTAATACGACTCACTATAGGGC |
| **Integration site 5' nested PCR** | Forward | AGACCCTTTTAGTCAGTGTGGAAAATC |
| | Reverse | ACTATAGGGCACGCGTGGT |

#### Supplementary Figure 1

Gating strategy used in flow cytometric analysis experiments as shown in Figure 1F

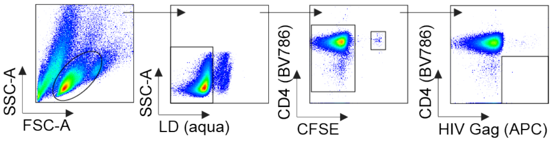

Gating strategy used in flow cytometric analysis experiments as shown in Figure 3D

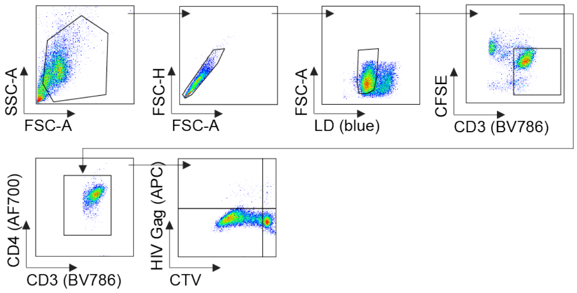

Gating strategy used in flow cytometric analysis experiments as shown in Figure 3B

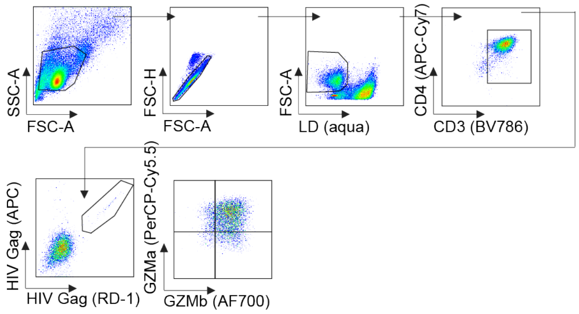

Gating strategy used in flow cytometric analysis experiments as shown in Figure 4E dARC COG5 and Figure 4G

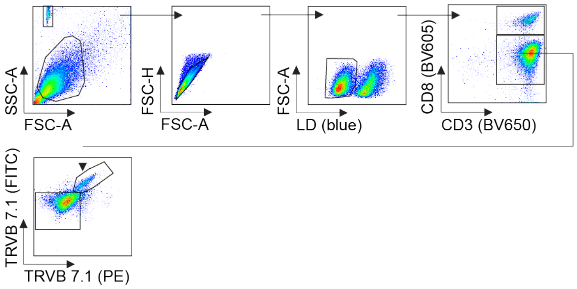

### Supplementary Figure 1

Gating strategy used in flow cytometric analysis experiments as shown in Figure 4A

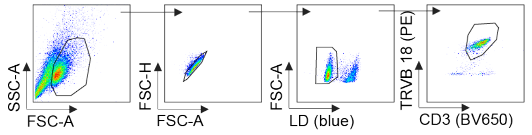

Gating strategy used in flow cytometric analysis experiments as shown in Figure 4E ARC Chr16 and Figure 4G

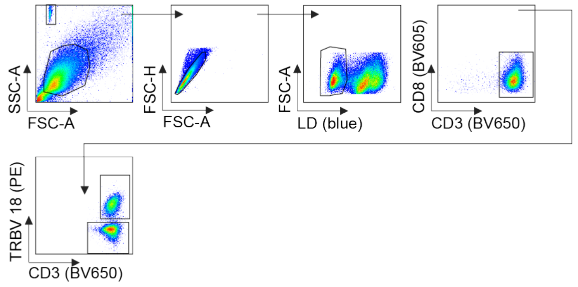

Gating strategy used in flow cytometric analysis experiments as shown in Figures 5F

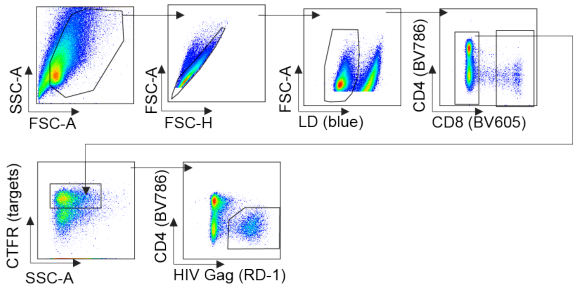

Gating strategy used in flow cytometric analysis experiments as shown in Figures 5H

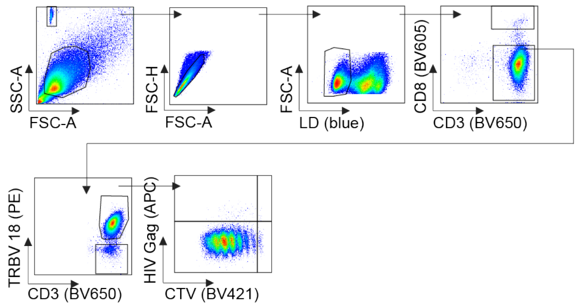
